# From unsupervised clustering to atlas-guided annotation in cohort-scale spatial omics with HiCAT

**DOI:** 10.64898/2026.05.27.728266

**Authors:** Jing Huang, Xueqi Shen, Yoland Smith, Lara Harik, Linghua Wang, Jindan Yu, Michael P. Epstein, Jian Hu

## Abstract

Pathologist-annotated tissue regions provide a fundamental reference for examining spatial omics data, yet such annotations are available for a limited number of samples due to the substantial manual effort required. Moreover, these annotations are derived from morphology within individual histology images, which can overlook molecularly defined regions and obscure intra-sample heterogeneity. To address these limitations, we present HiCAT, a machine-learning framework that automatically generates pathologist-informed region annotations and characterizes regional heterogeneity in spatial omics data. Across seven datasets, HiCAT consistently outperforms state-of-the-art methods, achieving a median relative improvement of 107% in accuracy. Beyond transferring pathologist annotations, HiCAT uncovers molecularly informed regional heterogeneity not captured by original annotations, including tumor subregions associated with clinical outcomes and brain subregions aligned with spatiotemporal disease progression. By generating consistent, highly granular, and biologically informative region annotations across large cohorts, HiCAT enables scalable downstream analysis and provides training labels for foundation models in spatial biology.

## Introduction

Spatial omics represents a paradigm shift in the study of complex biological systems by enabling molecular profiling while preserving the spatial context of tissue architecture^1^. Among spatial omics technologies, spatial transcriptomics (ST) is particularly powerful yet analytically demanding because of its high dimensionality and frequent integration with complementary modalities such as histology and protein measurements. As ST becomes more accessible, studies are increasingly generating multi-sample datasets spanning disease conditions and developmental stages^2-5^. The transition from single-sample to cohort-scale designs enables systematic comparisons across patient groups and longitudinal trajectories, creating new opportunities to uncover disease mechanisms and strengthen the generalizability of spatial biology findings.

Pathologist-annotated tissue regions provide a fundamental reference for interpreting spatial omics data. Compared with regions derived from unsupervised clustering, expert-defined regions are typically more biologically meaningful and clinically relevant^6^. However, such annotations typically are available for only a small number of sections and, as ST datasets scale to cohorts, manually annotating every section becomes laborious and impractical. This lack of expert supervision also constrains the recent development of foundation models (FMs) in genomics^7-9^. Unlike many pathology FMs^10,11^, where large expert-labeled corpora support supervised pretraining, most genomics FMs are trained in an unsupervised manner, sometimes yielding inferior performance to regression baseline in certain tasks^12,13^. Together, these gaps motivate supervised approaches that can propagate region annotations from labeled to unlabeled sections, enabling scalable construction of annotated atlases for cohort-level analysis.

Several state-of-the-art methods have been developed for supervised domain detection in ST. For example, STELLA^14^ uses graph convolutional networks to learn latent representations for label transfer from reference to query samples; SCGP^15^ applies Leiden community detection on spatial graphs for label propagation; SpaDo^16^ transfers annotations by matching query to reference embeddings derived from cell type and local niche composition. Although these approaches show promising performance, they fall short of two requirements in cohort-scale studies. First, most methods operate in a pairwise transfer setting, training on a single annotated section to label one unannotated section at a time, which limits their ability to learn and reconcile intra-sample heterogeneity in complex tissues. Second, their learning signals are largely constrained to morphology-guided regions derived from expert annotation. In practice, many biologically important tissue states are molecularly distinct yet morphologically similar and are therefore under-annotated by pathologists^17,18^. Together, these limitations motivate methods that move beyond one-to-one label transfer and incorporate molecular signals to refine, discover, and harmonize region definitions across samples.

To address these gaps, we introduce HiCAT (**Hi**erarchical **C**ohort-scale **A**nnotation **T**ransfer for spatial omics), a supervised framework for cohort-scale region annotation in spatial omics. HiCAT integrates multi-modal inputs to learn region classifiers from multiple annotated references, enabling accurate label transfer to large collections of unlabeled sections. Across seven datasets, HiCAT consistently outperforms existing methods, improving accuracy and F1 score. Beyond propagating original annotations, HiCAT can generate highly granular, disease-relevant maps in regions indistinguishable based on morphology alone. Together, HiCAT provides a scalable solution for building annotated spatial omics atlases and shifts cohort-level analysis from predominantly unsupervised workflows toward supervised, atlas-guided biological interpretation.

## Results

### Overview

As shown in **Fig. 1**, HiCAT is designed to integrate multi-modal spatial inputs, such as gene expression, protein abundance, and histology. In Step 1, HiCAT learns from multiple annotated references to derive a tissue hierarchical tree. This tree aggregates all tissue regions across samples, including regions that absent in individual sections. The hierarchy integrates morphology, spatial neighborhood, and molecular similarity to enable coarse-to-fine annotation, allowing different features to be emphasized at different branches and improving accuracy over one-step annotation. Motivated by the observation that normal regions (e.g., connective tissue) are relatively conserved across samples whereas disease-associated regions (e.g., tumor) are more variable^19,20^, HiCAT defines a heterogeneity score for each leaf in the tree to quantify cross-sample variability and prioritize potentially disease-relevant regions for further investigation. In Step 2, HiCAT compares each unannotated sample with the reference pool, excludes outlier references, and transfers labels using only the most compatible samples to avoid performance degradation from irrelevant references. In Step 3, HiCAT detects anchors using two complementary strategies. For within-study transfer, a k-nearest-neighbor (kNN)-based framework identifies anchors for each binary reference node using Euclidean distances in the selected feature space. For cross-study transfer, where domain shifts may occur, a quantile-based framework detects anchors using expression-level quantiles to ensure robustness. Together, these strategies allow HiCAT to adapt to different transfer scenarios and support reliable tissue-region annotation. In the Step 4, HiCAT performs label transfer and, for regions with high heterogeneity scores, supports automatic subtyping to resolve intra-sample spatial patterns beyond the granularity of the original annotations. The resulting annotated atlas enables cohort-level comparisons, such as contrasting healthy versus diseased samples to identify region-specific disease-associated features and analyzing time-series spatial omics datasets to characterize spatiotemporal changes during disease progression.

**Fig. 1.**
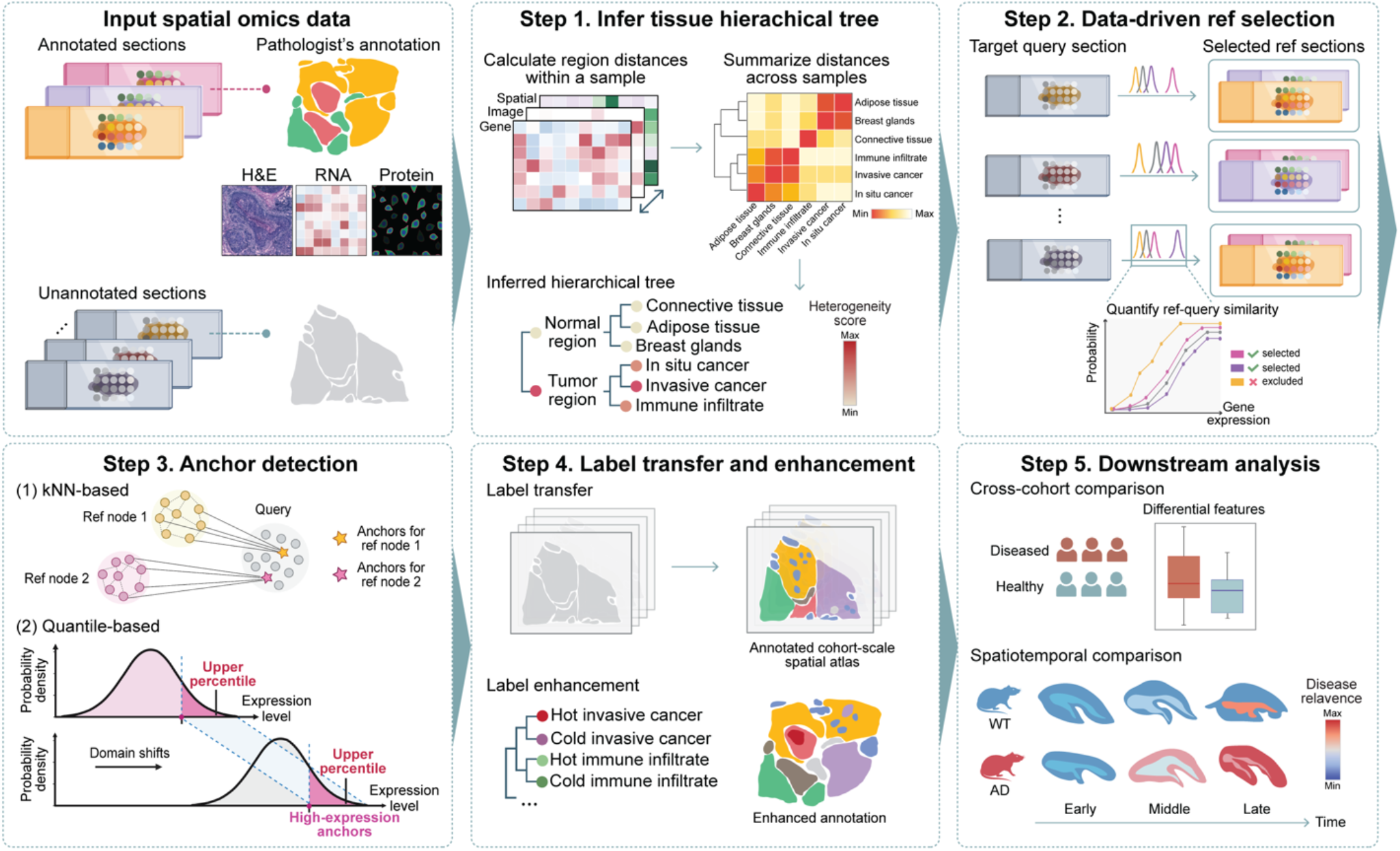
Workflow of HiCAT. HiCAT takes multi-modal spatial omics data as input, including paired H&E images, gene expression, and protein abundance, together with pathologist-provided scribble annotations to guide label transfer. In step 1, HiCAT learns from annotated reference samples to construct a hierarchical tree that captures tissue region organization. This tree is inferred by integrating molecular, morphological, and spatial neighborhood composition information to guide coarse-to-fine annotation. Region-level heterogeneity level is quantified across samples to prioritize disease-relevant regions. In step 2, for a given query sample, HiCAT evaluates its similarity to the reference pool using gene expression distributions and selects a subset of references for matched supervision. In step 3, HiCAT identifies anchors for each binary reference node using a kNN-based framework for within-study transfer or a quantile-based framework for cross-study or cross-technology transfer, where anchors are detected using upper expression quantiles rather than absolute expression values. In step 4, HiCAT performs hierarchical label transfer and, in regions with high heterogeneity, further resolves fine-grained subtypes beyond the original annotations. The resulting annotated sections enable construction of a spatial atlas for cohort-level comparisons between health and diseased samples and for characterizing spatiotemporal changes during disease progression.

### Benchmarking on breast cancer datasets

To evaluate the performance of HiCAT, we systematically compare it with four state-of-the-art label-transfer approaches^14-16,21^ using an HER2-positive breast cancer dataset^2^. This dataset includes 36 slides from eight patients (A–H), with one pathologist-annotated slide per patient serving as the ground truth. We selected the annotated sample of H1, G2, and E1 as the training samples as they encompass a wide diversity of tissue regions (**Fig. 2a**). The remaining five sections (A1, B1, C1, D1, and F1) were reserved as testing set (**Fig. 2b**). We use accuracy and F1 score as the evaluation metrics. For a fair comparison, methods capable of leveraging multiple samples for training (HiCAT, SCGP^15^, and Seurat^21^) were trained jointly on all three reference samples, whereas methods restricted to single-sample training (SpaDo^16^ and STELLAR^14^) were trained separately on each reference sample, and their best performance was reported. Detailed information on the training reference samples used by each method is provided in **Supplementary Table 1**. As summarized in **Fig. 2c**, HiCAT consistently achieves superior performance across all five test samples (median accuracy = 0.74, median F1 = 0.80), yielding a 68%–335% improvement in accuracy and a 78%–300% improvement in F1 score relative to other methods. **Fig. 2d** presents detailed results for all methods. HiCAT’s superior performance stems from its ability to accurately delineate major tissue regions, such as normal versus tumor region, and further distinguish tumor subtypes, including *in situ* cancer and invasive cancer. In contrast, SCGP fails to separate normal and tumor regions across all samples; STELLAR incorrectly assigns large tissue areas as unrecognized (grey); Seurat and SpaDo fail to distinguish *in situ* (orange) from invasive cancer (red), particularly in samples D1 and F1, and misclassify connective tissue (green) as immune infiltration (pink) in samples B1, C1, and D1.

**Fig. 2.**
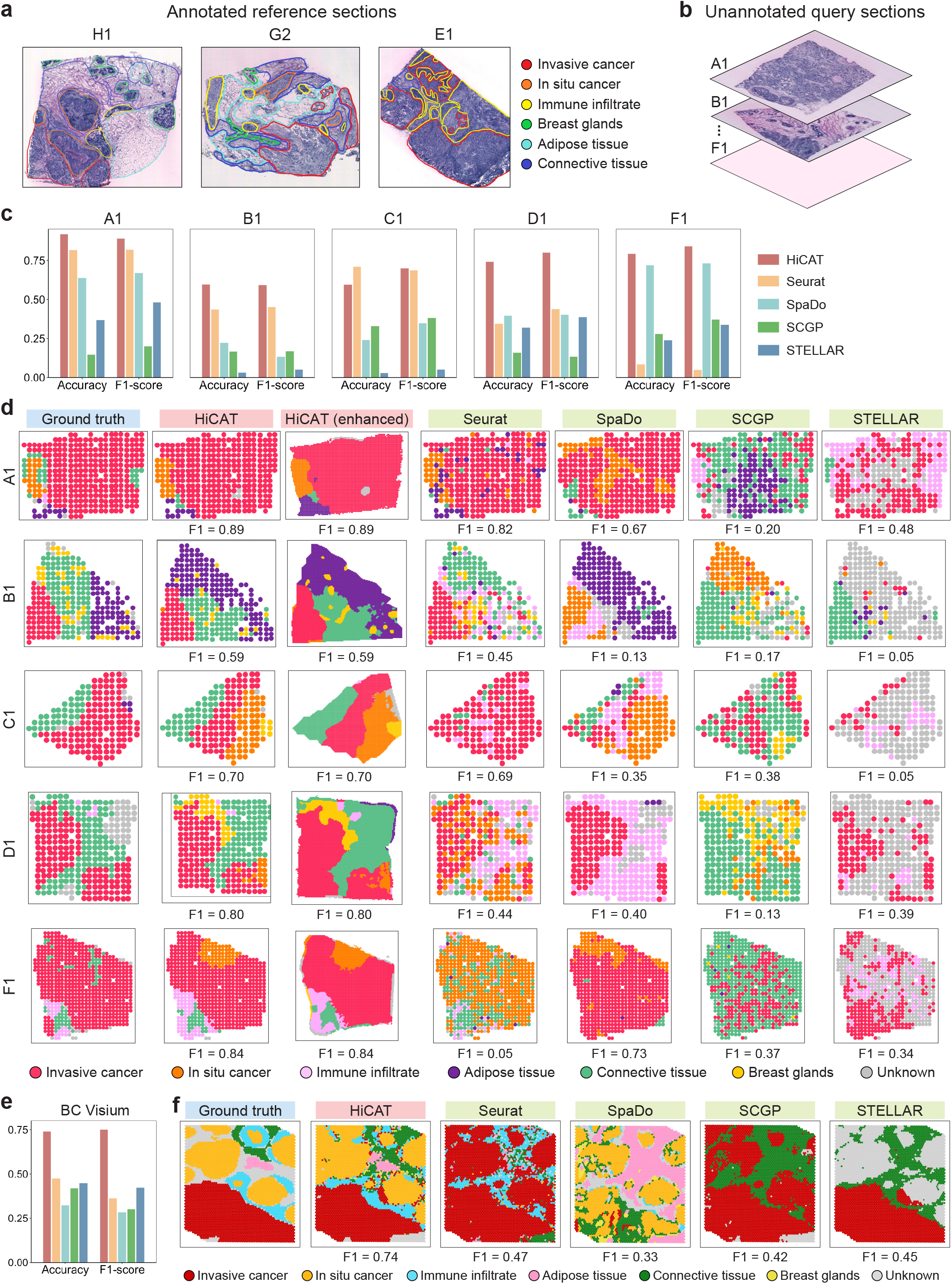
Benchmarking label transfer performance in breast cancer datasets. **a**. Annotated reference sections from HER2-positive breast cancer patients (H1, G2, E1), with pathologist-generated scribble annotations overlaid on H&E images. **b**. Unannotated query sections from HER2-positive breast cancer patients (A1, B1, C1, D1, F1) and a 10x Visium breast cancer sample from an independent study used for label transfer evaluation. **c**. Bar plots comparing within-study supervised label transfer performance across methods on HER2-positive query sections, evaluated using accuracy and F1 score. **d**. Tissue region annotation results on HER2-positive query sections. HiCAT provides both spot-level and enhanced-resolution region delineation, and other methods are shown using their best-performing results for comparison. **e**. Bar plots comparing cross-study supervised label transfer performance from HER2-positive reference sections to an independent 10x Visium breast cancer sample, evaluated using accuracy and F1 score. **f**. Tissue region annotation results on the 10x Visium query sample.

Beyond the major regions that dominate overall accuracy and F1 scores, HiCAT more effectively captures fine-grained, biologically important structures. For example, in sample B1 (**Fig. 2d**), HiCAT detects breast gland islands (yellow) with a precision of 0.81, exceeding the second-best method, Seurat (prevision=0.63). In sample D1, HiCAT identifies immune-cell infiltration (pink) with a precision of 0.95, outperforming Seurat (prevision=0.06). A detailed region-level evaluation is provided in **Supplementary Note 1**. Notably, these structures occupy only a small fraction of the tissue and have coarse boundaries at the native spot resolution (55-µm radius). By enabling coarse-to-fine annotation, HiCAT improves boundary delineation and accuracy (**Fig. 2d; HiCAT enhanced**).

In addition to comparisons with supervised methods, we also benchmarked HiCAT against unsupervised clustering approaches, including IRIS^22^, SCGP (unsupervised version), Seurat (unsupervised version), BayesSpace^23^, BASS^24^, and STAGATE^25^. HiCAT achieved the best overall performance, with a mean ARI of 0.44, representing a 29%–340% improvement over the comparison methods (**Supplementary Note 2**).

Next, we evaluated a more challenging scenario in which query samples were generated in different studies. Such cross-study data often exhibit strong batch effects, making robustness to this variability essential for cohort-scale analysis. Using the same evaluation framework, we transferred labels from HER2-positive ST samples to a 10x Visium breast cancer sample from an independent study^26^. Detailed information on the training references for each method is summarized in **Supplementary Table 2**. As shown in **Fig. 2e**, HiCAT achieved superior performance, reaching an accuracy of 0.82 and an F1-score of 0.84, corresponding to 74% and 133% improvements over the second-best method, respectively. The detailed annotations in **Fig. 2f** show that HiCAT accurately separates *in situ* cancer from invasive cancer, whereas Seurat, SpaDo, and SCGP fail to distinguish these regions. Notably, adipose tissue (pink) and connective tissue (green) occupy only a small fraction of this test sample, and only HiCAT successfully identifies them while other methods fail (**Fig. 2f**). Compared with the second-best method Seurat, HiCAT improved precision by 0.4 and 0.17 for adipose and connective tissue, respectively (**Supplementary Fig. 3**). In addition, comparisons with six unsupervised methods further highlight the superior performance of HiCAT, as detailed in **Supplementary Note 3**. Together, these results demonstrate HiCAT’s ability to perform robust cross-study label transfer and establish a foundation for cohort-level ST data integration and analysis.

### HiCAT characterizes cross-sample heterogeneity in breast cancer

The heterogeneity scores on the tree in **Fig. 3a** highlight *in situ* cancer, invasive cancer, and immune infiltration as the most variable regions across samples. HiCAT further resolves fine-grained subtypes within these regions (**Fig. 3b**) that are not captured by the original annotations in the three reference samples. Next, HiCAT interprets heterogeneous subtypes using differential expression (DE) and Gene Set Enrichment Analysis (GSEA). As shown in **Fig. 3c**, both *in situ* and invasive cancer harbored a subtype present across all three samples, characterized by MHC class I-mediated antigen presentation and T cell cytotoxicity genes (*HLA-A/B/C, B2M*) together with type I interferon signaling markers (*IFI27, STAT1*), consistent with an immune-inflamed, “hot” tumor state^27,28^. By contrast, the remaining cancer regions exhibited comparatively immune-cold profiles. Within immune regions, HiCAT identified subtypes enriched for antigen processing and presentation through MHC class I and II pathways (*HLA-B, B2M, HLA-DRA*)^29^ and iron regulation (FTL)^30^, indicating an activated immune niche (**Fig. 3d**). In the zoomed-in views of samples G2 and E1 (**Fig. 3b**), activated immune cells exhibit a shared spatial pattern, concentrating at the boundary of the immune aggregate closest to the hot cancer region, whereas non-activated immune cells are distributed farther from the tumor. These finely resolved subtypes may appear as sparse points at the original resolution, but HiCAT’s enhanced-resolution predictions enable detailed subtype-level examination beyond the native spot resolution. To quantitatively assess differences in spatial neighborhood relationships among these subtypes, we calculated inter-region distances (**Fig. 3e**) and found that hot tumor subtypes were located closer to activated immune subregions (mean distance=0.38) than cold tumor subtypes (mean distance=0.59), forming discrete tumor–immune foci. These patterns were consistently observed across the remaining five test samples (**Supplementary Note 4**), highlighting tumor–immune niches revealed by HiCAT’s heterogeneity analysis beyond label transfer.

**Fig. 3.**
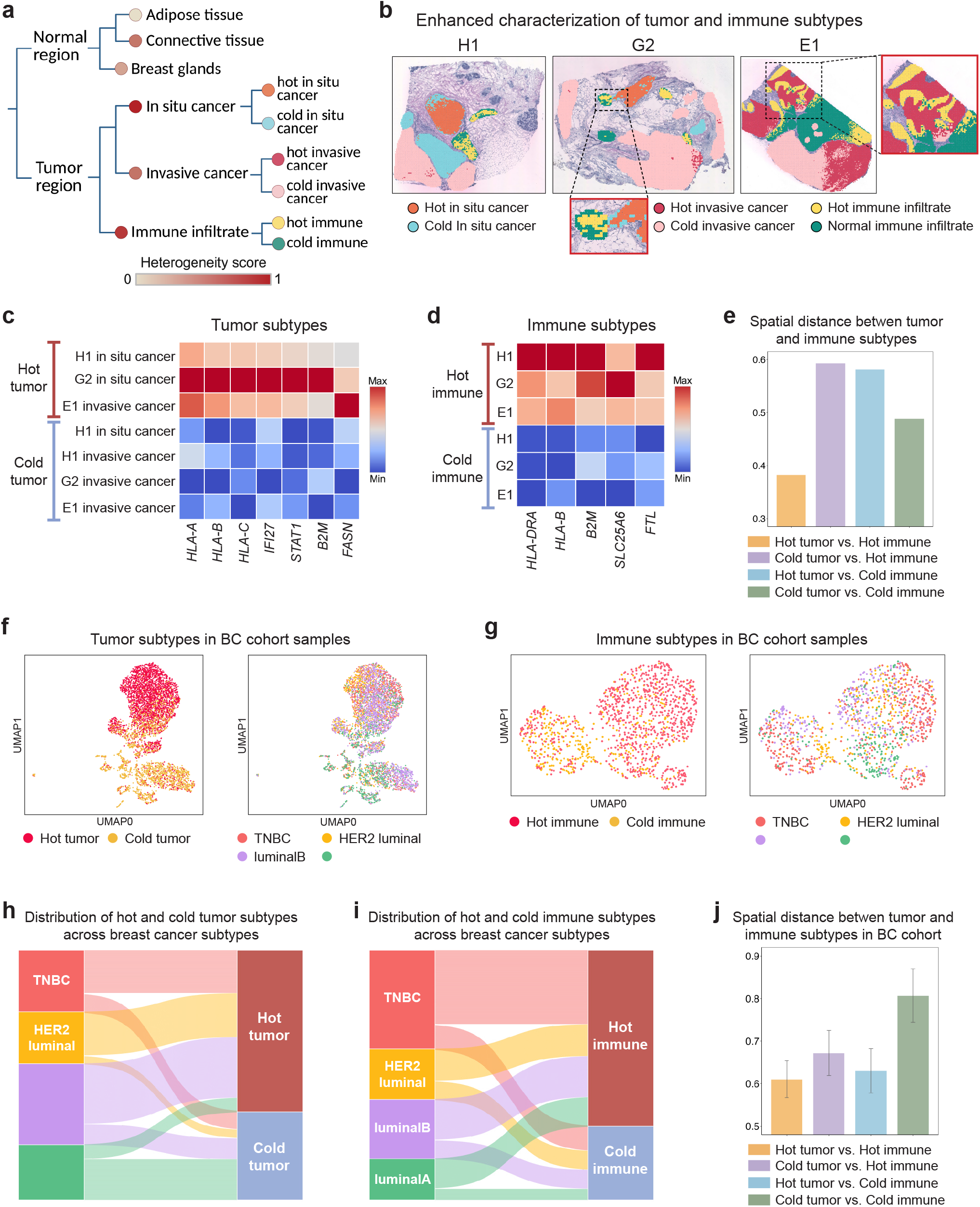
Characterization of cross-sample heterogeneity in cohort-level breast cancer datasets. **a**. Hierarchical tree inferred from HER2-positive breast cancer reference sections, capturing tissue region organization in breast cancer. Node color indicates the heterogeneity score of each region. For heterogeneous regions, HiCAT further resolves fine-grained subtypes, with node color indicating subtype identities (consistent with **b**). **b**. Enhanced-resolution annotation plots depicting the spatial distribution of fine-grained subtypes within heterogeneous regions. Zoom-in views highlight tumor-immune niches, with activated immune cells localized near or embedded within hot tumor regions. **c-d**. Gene expression enrichment heatmaps illustrating shared molecular programs across reference sections. **c**. Immune-inflamed, “hot” tumor state. **d**. Activated, “hot” immune niche. **e**. Spatial distance between tumor and immune subtype regions in HER2-positive reference sections, with bar heights representing mean distances computed across spots and sections. **f-g**. Cohort-level embedding of tumor and immune heterogeneity. **f**. UMAP visualization of tumor spots, colored by tumor subtype (hot vs. cold; left) and breast cancer clinical subtypes (TNBC, HER2 luminal, luminalB, and luminalA; right). **g**. UMAP visualization of immune spots, colored by immune subtype (hot vs. cold; left) and clinical subtype (right). **h-i**. Distribution of molecular subtypes across clinical subtypes. Sankey plot of tumor (**h**) and immune (**i**) subtypes across breast cancer clinical subtypes. **j**. Spatial distance between tumor and immune subtype regions across the breast cancer cohort. Bar heights represent mean distances across spots and sections, with error bars indicating standard error.

To demonstrate the scalability of HiCAT for cohort-level analysis, we applied it to a breast cancer ST cohort comprising 18 patients across four clinical subtypes^31^: luminal A (n = 4), luminal B (n = 5), HER2 luminal (n = 5), and TNBC (n = 4). Using HER2-positive ST samples as references, HiCAT automatically selected suitable references for each test sample (**Supplementary Table 3**), transferred region labels to all 18 sections (**Supplementary Fig. 6**), and characterized tumor and immune heterogeneity. Across patients, HiCAT consistently recovered hot and cold tumor subtypes (**Fig. 3f**) as well as activated and non-activated immune subtypes (**Fig. 3g**), as shown in the UMAP embedding. The identities of these subtypes were further supported by enriched marker genes and pathways (**Supplementary Fig. 7**). The Sankey plot in **Fig. 3h** revealed that TNBC, HER2-luminal, and luminal B contained a higher proportion of hot tumor regions (70%, 84%, and 74%, respectively) than luminal A (26%). This pattern is consistent with the literature showing that luminal A breast cancer is a lower-proliferation subtype than the others^32,33^. Further examination of proliferation-related genes, including *PCNA, TOP2A, and CDC20* (**Supplementary Fig 8**), also supported this observation. In contrast, the proportions of immune subtypes were broadly similar across different subtypes (**Fig. 3i**). Distance analysis showed that, across all subtypes, hot tumor regions were consistently closer to activated immune subtypes than distances among other subtype pairs (**Fig. 3j**, one-side two-sample t-test p=0.047), highlighting a conserved tumor–immune interaction pattern. Together, these results demonstrate HiCAT’s ability to uncover clinically relevant insights from cohort-scale spatial transcriptomics analysis.

### Benchmarking on Multi-omics Tonsil

As spatial technologies expand into multi-omics, analysis methods increasingly need to handle multi-modal inputs. Although different modalities can provide complementary information, they can also introduce conflicting signals because each modality differs in resolution, noise characteristics, and the biological processes it captures. As a result, directly integrating additional modalities does not always improve performance and can even reduce domain detection accuracy, as reported for several multi-omics integration methods^18,34^. A desirable property of multimodal spatial analysis is therefore selective integration, so that complementary signals are retained while conflicting or noisy information is minimized.

To demonstrate that HiCAT has this desirable property, we benchmarked it against competing methods using a tonsil spatial multi-omics dataset comprising two tissue sections with simultaneous profiling of H&E morphology, gene expression, and protein abundance^35^. As shown in **Fig. 4a**, our pathologist annotations were used as ground truth, and labels were transferred from section 1 (**Fig. 4a**) to section 2 (**Fig. 4b**). As shown in **Fig. 4c** and **4d**, HiCAT consistently benefited from adding additional modalities. Starting with gene expression alone and then incorporating morphology and protein information, HiCAT’s F1 score increased from 0.72 to 0.74 to 0.77, respectively. By contrast, other methods showed limited or inconsistent gains from multimodal input. Seurat achieved an accuracy of 0.68 regardless of whether protein data were included, SCGP decreased from 0.54 to 0.49 after adding protein data, and STELLAR improved from 0.13 to 0.22 but remained the lowest-performing method overall. Accuracy in **Fig. 4d** showed the same trend.

**Fig. 4.**
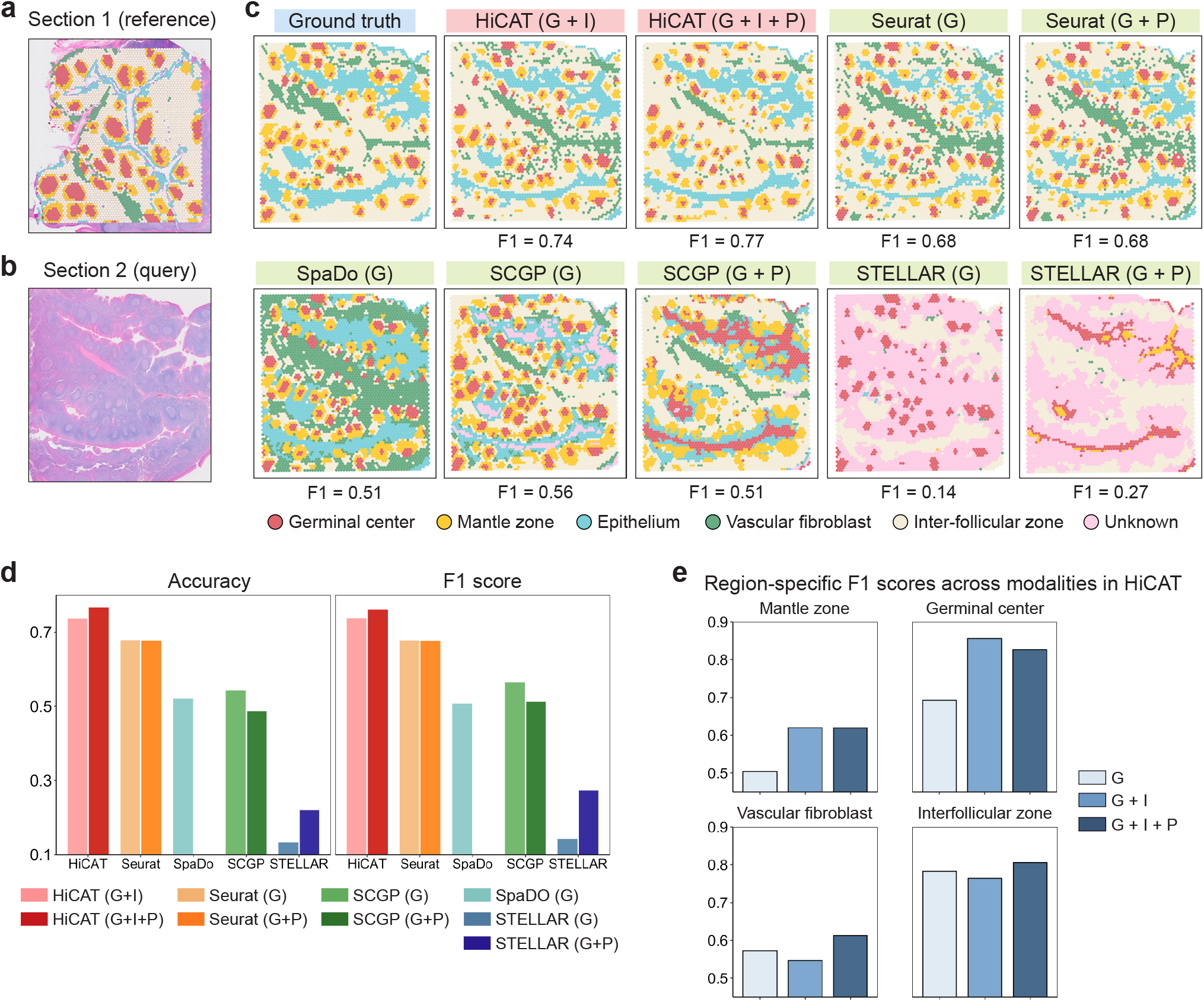
Benchmarking label transfer performance in a multi-omics tonsil dataset. **a**. Annotated reference section of tonsil Visium Section 1, with pathologist-generated annotations overlaid on the H&E image. **b**. Unannotated query section of tonsil Visium Section 2. **c**. Tissue region annotation results for the query section. HiCAT results are shown using two modalities (RNA and histology) and three modalities (RNA, histology, and protein). Seurat, SCGP, and STELLAR results are shown using RNA alone and two modalities (RNA and protein). SpaDo results are shown using RNA alone. **d**. Bar plots comparing supervised label transfer performance across methods and modality settings on the tonsil spatial omics query sample, evaluated using accuracy (left) and F1 score (right). **e**. Bar plots comparing region-specific F1 scores under different modality settings in HiCAT, highlighting the contribution of each modality to region-level detection.

Notably, although the overall accuracy gain from incorporating additional modalities into HiCAT was modest, further analysis showed that these modalities substantially improved the identification of specific fine-grained regions rather than uniformly enhancing overall performance. Specifically, image features most strongly improved the detection of the germinal center and mantle zone, likely because these regions have distinctive morphology. As shown in **Fig. 4e**, adding H&E information to gene expression in HiCAT improved the mantle zone F1 score from 0.50 to 0.62 and the germinal center F1 score from 0.69 to 0.86. Although these regions occupy only a small fraction of the tissue and contribute little to overall accuracy, their precise delineation is important for interpreting tonsillar microarchitecture, B cell maturation, and immune regulation. Protein features, in turn, improved the detection of the vascular fibroblast and interfollicular zone. Furthermore, as shown in **Fig. 4e**, adding protein information to HiCAT increased the vascular fibroblast F1 score from 0.55 to 0.61 and the interfollicular zone F1 score from 0.76 to 0.81. Further examination of protein abundance patterns showed that only a subset of proteins provided strong discriminatory signal, whereas others contributed limited information. Specifically, PECAM1^36^ and ACTA2^37^, well-supported markers of vascular-associated regions, were informative for the vascular fibroblast region, while CD3E and CD8A, established T cell markers, distinguished the interfollicular zone^38^. These proteins were automatically selected by HiCAT to separate the corresponding regions (**Supplementary Fig. 9**). Notably, these region-specific gains from incorporating protein information were not observed in the competing methods (**Supplementary Fig. 10**). Consistent with its hierarchical annotation strategy, HiCAT was able to leverage informative features while down-weighting less informative signals in each modality, demonstrating its superior ability to selectively benefit from complementary multi-omics information.

### Application to Tonsil Visium HD

Emerging spatial platforms such as Visium HD are rapidly advancing toward higher resolution and larger gene panels, creating new challenges for method scalability and efficiency. To evaluate HiCAT on such data, we applied it to a tonsil Visium HD dataset at 8 μm × 8 μm resolution^39^. Using the spot-level Visium sample with 55 μm-diameter spots as the reference, we transferred labels to the Visium HD section (**Fig. 5a**). As shown in **Fig. 5b**, HiCAT achieved an accuracy and F1 score of 0.75, corresponding to 17%–316% higher accuracy and 25%–213% higher F1 score than competing methods. Beyond overall accuracy, HiCAT consistently achieved the highest region-specific F1 scores in **Fig. 5c**. Notably, HiCAT was the only method that precisely delineates the small but biologically important immune structures of the germinal center (red) and mantle zone (yellow) within the secondary lymphoid follicle (**Fig 5a**). In addition to superior performance, HiCAT is also computationally efficient, achieving an average runtime of 84s and memory usage of 3.9GB across ten independent runs—approximately one-eighth the runtime and one-seventh the memory of competing methods as shown in **Fig. 5d** (SpaDo failed to process this dataset due to memory constrains).

**Fig. 5.**
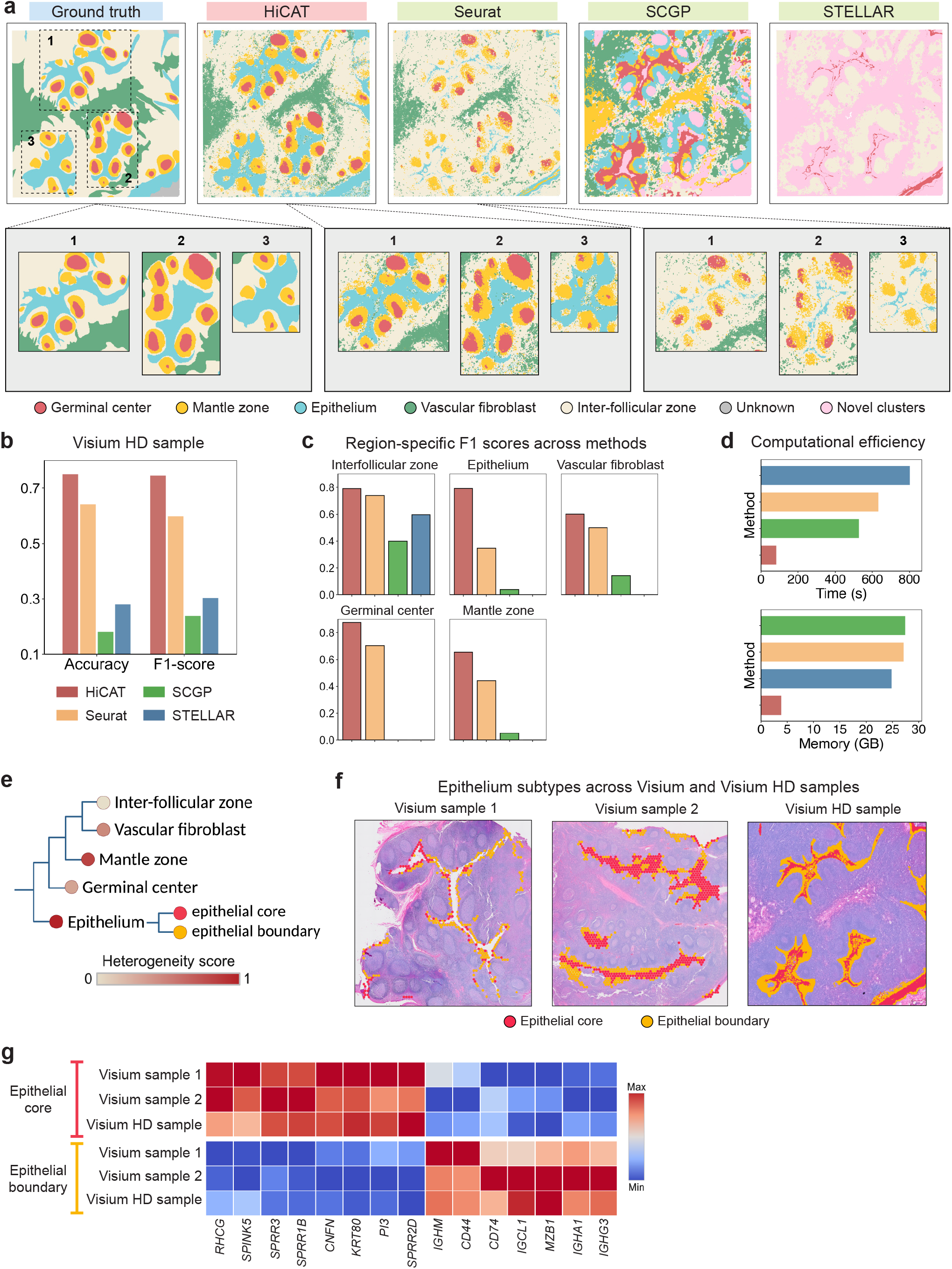
Benchmarking label transfer performance in a Visium HD tonsil dataset. **a**. Tissue region annotation results transferring labels from Visium reference tonsil samples to a Visium HD query sample. Zoom-in views highlight the delineation of secondary lymphoid follicle structures, comparing ground truth with HiCAT and the second-best-performing method of Seurat. **b**. Bar plots comparing supervised label transfer performance across methods on the Visium HD sample, evaluated by accuracy and F1 score. **c**. Bar plots comparing region-specific F1 scores across methods. **d**. Bar plots comparing runtime and memory usage across methods on the Visium HD sample. Bar heights indicate the average runtime and memory usage across ten independent runs. **e**. Hierarchical tree inferred from Visium reference tonsil sections, capturing tissue region organization in tonsil. Node color indicates the heterogeneity score of each region. For heterogeneous regions, HiCAT further resolves fine-grained subtypes, with node color indicating subtype identities (consistent with **f**). **f**. Epithelium subtypes identified across Visium and Visium HD samples, overlaid on corresponding H&E images. **g**. Gene expression enrichment heatmap illustrating shared molecular programs of epithelium subtypes across Visium and Visium HD samples.

**Fig. 5e** shows that the epithelial region displayed the greatest heterogeneity across the two Visium and one Visium HD samples. HiCAT consistently identified two epithelial subtypes with distinct spatial organization (**Fig. 5f**) and expression (**Fig. 5g**), forming an inner-core versus outer-boundary pattern. The boundary subtype, located at the interface with the interfollicular zone, was enriched for immunoglobulin genes (*IGHA1, IGHG3, IGLC1, IGHM*) and plasma cell-associated markers (*MZB1, CD74*), supporting active humoral immune engagement and antigen-associated processes at the mucosal interface^40,41^. In contrast, the inner subtype showed elevated expression of keratinization genes (*KRT80, PI3*), epithelial differentiation markers (*CNFN, RHCG*), and barrier-related genes (*SPRR3, SPRR1B, SPRR2D, SPINK5*), suggesting a differentiated core that maintains epithelial barrier integrity^42^. Together, these findings provide insight into epithelial–immune crosstalk within the tonsillar mucosal microenvironment.

### Application to AD and WT time series mouse brain data

A growing trend in recent studies is to generate spatial omics data along disease trajectories to analyze spatiotemporal dynamics. To demonstrate HiCAT’s capability to analyze such data, we applied it to a 10x Visium mouse brain dataset^43^. As shown in **Fig. 6a**, this dataset comprises two biological replicates per time point from TgCRND8 transgenic mice sampled at 2.5, 5.7, and 17.9 months (n=6), and wild-type littermates sampled at 2.5, 5.7, and 13.2 months (n=6). We used one sample per time point/condition for training and evaluated on the remaining six samples using neuropathologist annotations as ground truth. For all competing methods, we used the paired biological replicate as reference for each test sample, as samples matched by time point and genotype are most similar. The specific reference samples used by each method are provided in **Supplementary Table 4**. As shown in **Fig. 6b**, HiCAT consistently shows superior performances across six test samples (median accuracy = 0.83, median F1 score = 0.83), yielding an 8%-246% improvement in accuracy and 6%-301% improvement in F1 score compared to other methods. Detailed annotation plots are shown in **Supplementary Fig. 11**. Further examination showed that HiCAT also substantially improved the delineation of key regions, including the cornu ammonis (CA) and dentate gyrus (DG) within the hippocampal formation (HPF), as well as the intricate cortical layers of the isocortex (**Supplementary Note 5**).

**Fig. 6.**
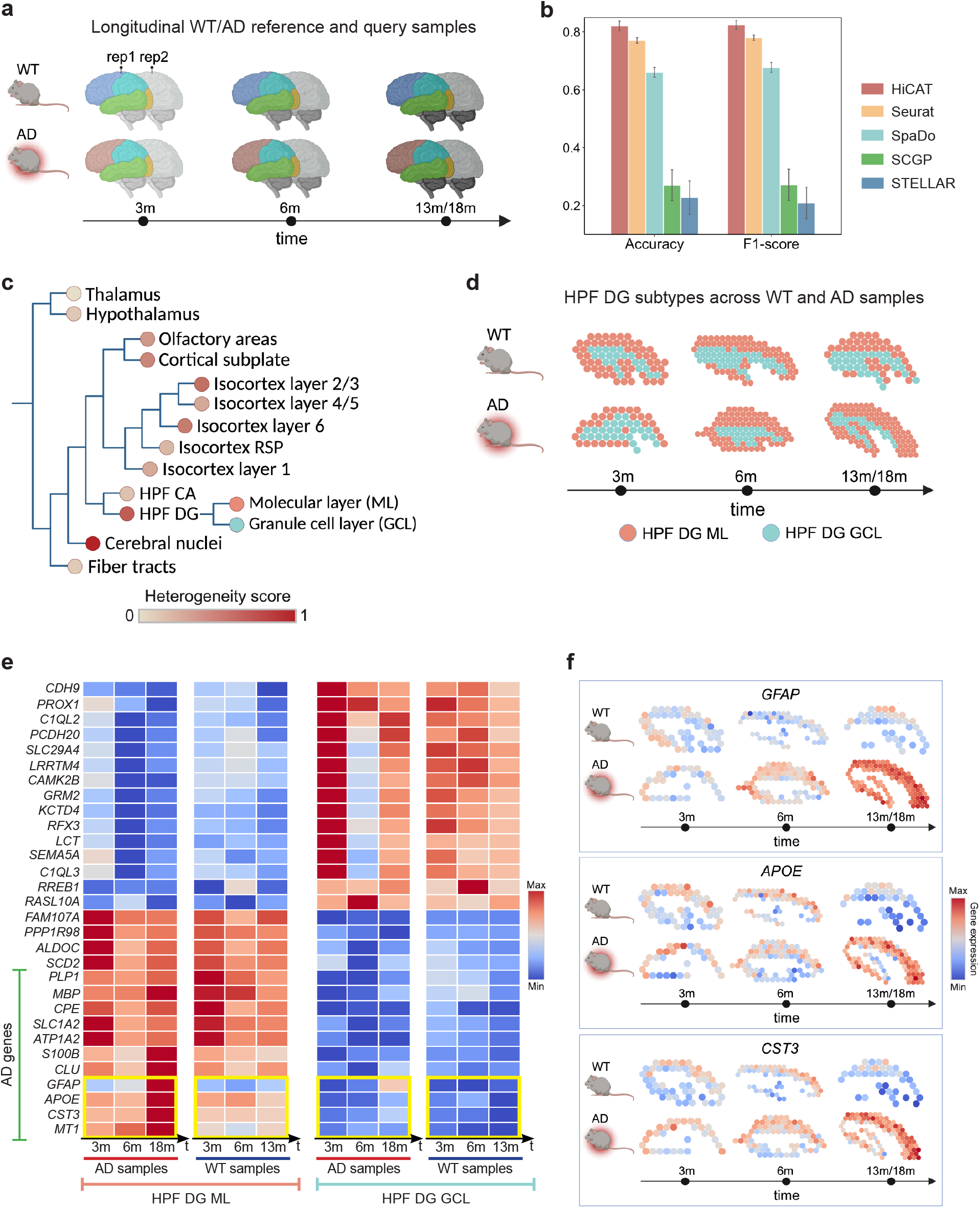
Benchmarking and heterogeneity characterization in a spatiotemporal mouse brain dataset across disease conditions. **a**. Study design of the mouse brain dataset. The dataset includes two biological replicates per time point, comprising transgenic AD mice at 2.5, 5.7, and 17.9 months (*n* = 6) and wild-type (WT) littermates at 2.5, 5.7, and 13.2 months (*n* = 6). One sample per time point and condition is used as reference, and the remaining six samples are used as queries. **b**. Bar plots comparing supervised label transfer performance across methods on mouse brain query samples. Bar heights indicate mean accuracy and F1 score across query samples, with error bars representing standard error. **c**. Hierarchical tree inferred from mouse brain reference samples, capturing tissue region organization in brain. Node color indicates the heterogeneity score of each region. For heterogeneous regions, HiCAT further resolves fine-grained subtypes, with node color indicating subtype identities (consistent with **d**). HPF CA denotes hippocampal formation Ammon’s horn, and HPF DG denotes hippocampal formation dentate gyrus. **d**. HPF DG subtypes identified across mouse brain reference samples, spanning disease conditions and time points. ML denotes molecular layer, and GCL stands for granule cell layer. **e**. Gene expression enrichment heatmap illustrating shared molecular programs of HPF DG subtypes across reference and query samples. Nine of the top 15 genes enriched in HPF DG molecular layer are known AD-associated marker genes (green lines). Among these AD risk genes, four exhibit progressive increases over time specifically in AD samples (highlighted by the yellow rectangle). **f**. Gene expression patterns of AD risk genes within the HPF DG molecular layer region in reference samples across disease conditions and time points.

Next, we investigated heterogeneity to characterize region-specific dynamics during AD progression. The hierarchical tree in **Fig. 6c** recapitulated the major anatomical organization of the brain, separating structures such as the layered isocortex and the HPF into distinct CA and DG compartments. HiCAT identified HPF-DG and the cerebral nuclei as the most heterogeneous regions. We focused on HPF-DG given its central role in memory and cognition. Beyond the original annotation, HiCAT further resolved HPF-DG into two subtypes (**Fig. 6d**), corresponding to the molecular layer (ML) and granule cell layer (GCL). DE analysis revealed distinct molecular profiles for these subtypes (**Fig. 6e**). The GCL was enriched for genes associated with neuronal homeostasis and granule neuron identity, including synapse organization and maturation (*CAMK2B, LRRTM4, C1QL2, C1QL3, CDH9*), synaptic transmission (*GRM2, CAMK2B*), and granule neuron markers (*PROX1, CAMK2B*). In contrast, the ML showed increased expression of genes linked to AD-related processes, including amyloid–tau regulation and clearance (*APOE, CLU*), glial reactivity (*S100B, GFAP*), and stress-responsive proteostasis (*NDRG2, VIM, CST3*), with their gradient patterns over time in **Fig 6f**. Together, these results suggest that the ML is more vulnerable during AD progression, whereas the GCL remains relatively preserved, consistent with prior reports of selective susceptibility within hippocampal circuits^44-47^. This region-specific signal would be obscured if HPF-DG were analyzed as a single region under the original annotation, highlighting the importance of HiCAT for subregion-level annotation

## Discussion

In this work, we introduced HiCAT, a hierarchical annotation framework that propagates pathologist-defined regions while explicitly accounting for intra-sample heterogeneity. In addition to reproducing original morphology-guided labels, HiCAT refines region boundaries and identifies more granular subregions driven by molecular profiles, enabling more detailed atlas construction and downstream analysis.

We extensively validated HiCAT across diverse real-world datasets spanning tissue types, biological contexts, platforms, and assay designs, including human breast cancer, prostate cancer, tonsil, and a time-series mouse brain cohort. Across these settings, the hierarchical trees inferred by HiCAT captured biologically meaningful structures, such as the layered brain and cancer clones (**Supplementary Note 6**), providing a useful foundation for annotation. Compared with four supervised and six unsupervised methods, HiCAT more accurately recovered pathology-annotated regions across both dominant tissue domains and subtle yet biologically important structures. To demonstrate scalability, we applied HiCAT to a cohort-scale breast cancer dataset, where it revealed clinically relevant features beyond the original annotations. In the mouse brain time series, HiCAT further resolved spatiotemporal changes in region-specific programs during disease progression. HiCAT also demonstrated strong computational efficiency, scaling to Visium HD-level data while achieving more than 7-fold reduction in runtime and memory usage compared with alternative approaches. Notably, in a tri-modality tonsil dataset, HiCAT consistently benefited from additional modalities rather than suffering the performance degradation observed in several competing approaches. This performance-preserving integration is especially favorable as the field transitions from single-modality to multi-omics spatial profiling.

Despite these strengths, one remaining limitation for HiCAT is that heterogeneity subtypes discovered beyond the original annotations still require manual interpretation based on their enriched genes and pathways. One promising direction is to incorporate large language models to translate these gene and pathway signatures into concise, human-interpretable region identities.

In summary, HiCAT provides a scalable, flexible, and generalizable solution to key limitations in cohort-scale spatial omics. Without HiCAT, constructing region-annotated atlases at the scale increasingly produced by modern spatial platforms would require prohibitive pathologist effort and would remain bottlenecked by sparse, morphology-only labels. By addressing this gap, HiCAT reduces manual burden, improves cross-sample consistency, and enables systematic characterization of disease-relevant heterogeneity that is obscured in single-sample or purely unsupervised analyses. HiCAT also provides a practical path to richly labeled datasets that can support supervised, atlas-guided cohort comparisons and serve as training signals for next-generation foundation models in spatial biology.

## Acknowledgements

J.H. was supported by Emory AI Humanity Initiative, Georgia Clinical & Translational Science Alliance, and NIH R35GM159880. M.E. was supported by NIH RF1AG071170 and NIH R01MH126449.

## Author contributions

This study was conceived and led by J.Hu. and M.E.. J.Huang. designed the model and algorithm and implemented the HiCAT software with input from J.Hu. and M.E. J. Huang led data analyses with evaluation input and annotations from Y.S., L.H., L.W., and J.Y.. X.S. contributed to part of the data analysis. J.Hu. and J.Huang. wrote the paper with feedback from all other co-authors.

## Methods

### Preprocessing of spatial multi-modal inputs

HiCAT integrates multi-modal information from spatial omics data, including gene expression, protein abundance, and associated histology images. For gene expression, HiCAT takes the raw UMI count matrix as inputs and performs library-size normalization, followed by plus-one log transformation. For protein abundance, zero-clipping is applied, followed by normalization and plus-one log-transformation, to mitigate sparsity and technical dropouts. For the paired H&E image, HiCAT extracts image features using UNI^10^, a Vision Transformer (ViT)-based pathology foundation model. UNI takes 16×16-pixel patches as inputs and outputs a 2,048-dimensional feature vector for each patch. Each vector captures both local information within the 16×16-pixel patch and broader tissue context from the surrounding 224×224-pixel region centered on the patch. To align with spatial omics measurements, embeddings from all 16x16-pixel patches within each omics spot are averaged to obtain a spot-level image feature embedding. In addition to the default UNI, HiCAT also supports HIPT as an alternative, which utilizes a hierarchical ViT model to generate 576-dimensional embeddings and offers improved computational efficiency. For pathology scribble annotation, HiCAT uses an OpenCV-based contour detection pipeline to convert the annotations into spot-level labels.

### Problem formulation

We illustrate HiCAT using spatial data with two modalities: gene expression (*G*) and paired H&E-stained histology images (*I*). We note the framework can also incorporate additional modalities, such as protein abundance, which we illustrate using a tonsil multi-omics dataset. For each tissue section *s* with *n*_*S*_ spots, its corresponding feature matrix for each modality *m* is represented as 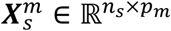, where *p*_*m*_ is the number of features for that modality. Let *S*_*ref*_ and *S*_*query*_ denote the sets of annotated reference sections and unannotated query sections, respectively. For each reference section *s* ∈ *S*_*ref*_, we have spot-level labels 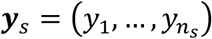, where *y*_*i*_ ∈ {1, …, *K*_*s*_} and *K*_*s*_ denotes the number of region categories for that section. HiCAT aims to leverage multimodal information from annotated references 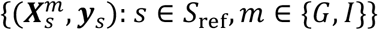 to infer region labels for spots in the unannotated query sections *S*_*query*_.

### Hierarchical Tree inference

HiCAT first derives a hierarchical tree to represent the relative similarity among tissue regions, which subsequently guides hierarchical label transfer. The hierarchy is inferred from all labeled reference sections *S*_*ref*_ incorporating omics measurements, morphology, and spatial proximity, as outlined in the following two steps:

#### Step 1: Calculating multi-modal pairwise distance

HiCAT first computes region similarity within each reference section by integrating modality-specific distances. These section-level similarities are then aggregated across all references to obtain a unified hierarchical relationship.

For each reference section *s* ∈ *S*_*ref*_ with modality *m* ∈ {*G, I*}, we identify region-specific features. For region *k* ∈ {1, …, *K*_*s*_}, its representative feature set 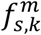is identified by DE analysis, comparing feature values in spots within versus outside the region *k* using the Wilcoxon rank-sum test. By default, features with FDR-adjusted P value < 0.05 and fold change > 1.15 are ranked by fold change, and the top 10 features are retained. A shared feature space is constructed by taking the union set of representative features across all regions, 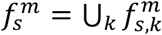 Each region is then represented by a centroid vector 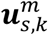, computed as the average feature values across its spots in the shared feature space 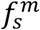

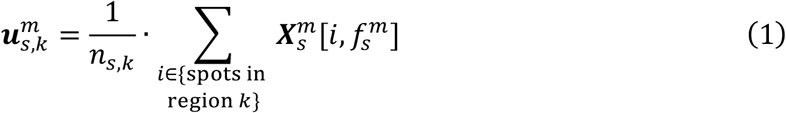

where *n*_*s,k*_ is the number of spots in region *k* of section *s* and 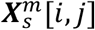 is the value of feature *j* at spot *i*. The pairwise distance between regions *k*_1_ and *k*_2_ is then quantified by the Euclidean distance between their centroid representations in this feature space:

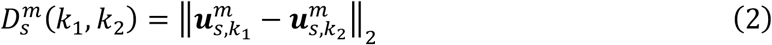

This yields modality-specific distances 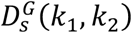 and 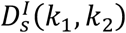, computed from gene expression and image features, respectively.

Regions that are spatially proximal often share similar microenvironments. To account for this, HiCAT incorporates spatial proximity when measuring regional similarity. We define a radius *r* such that each spot has an average of 5 neighbors within *r* . For any two regions *k*_1_, *k*_2_ ∈ {1, …, *K*_*s*_}, we aggregate the neighborhoods of all spots within *k*_1_ and compute the proportion of neighboring spots belonging to *k*_2_, denoted as 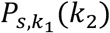. Regions that are spatially closer are expected to have higher mutual neighborhood proportions. We therefore define the spatial distance between regions as an inverse function of this mutual neighborhood composition:

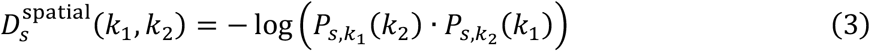

As distances from different modalities may have different scales, we apply min-max normalization to rescale them to range [0,1]. These normalized distances are then combined via a weighted sum to obtain a section-level integrated distance across modalities:

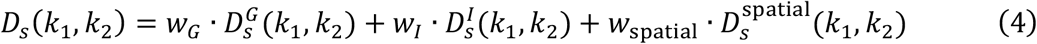

where *w*_*G*_ = *w*_*1*_ = *w*_spatial_= 1/3 by default.

To derive a unified distance measure across reference sections, HiCAT aggregates section-level distance across them. For a region pair *K*_1_,*K*_2_, let 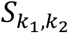 denote the set of reference sections that contain both regions. Distances are combined using a weighted average, with weights proportional to the number of spots in each section, given that larger sections tend to provide information:

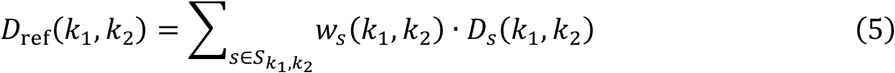

where *w*_*s*_(*k*_1_, *k*_2_) denotes the weight for section *s*. The resulting distance *D*_ref_(*k*_1_, *k*_2_) ∈ [0,1], with larger values indicate lower similarity.

#### Step 2: Tree construction

The hierarchical tree that captures tissue organization across samples is constructed based on the distance *D*_ref_(*k*_1_, *k*_2_) defined in Equation 5. HiCAT employs an agglomerative clustering with a greedy merging strategy to iteratively merge regions until all regions are merged into a single cluster, as detailed in the following pseudocode (Algorithm 1). The resulting hierarchy is represented as a binary tree structure *Tree* = (*V, E*), where *V* denotes cluster nodes and *E* encodes parent-child relationships. In this hierarchical tree, regions with similar multimodal profiles and spatial contexts cluster at lower levels, whereas higher levels capture broader tissue organization. This hierarchy provides a structured basis for subsequent label transfer.

##### Algorithm 1

for hierarchical tree inference

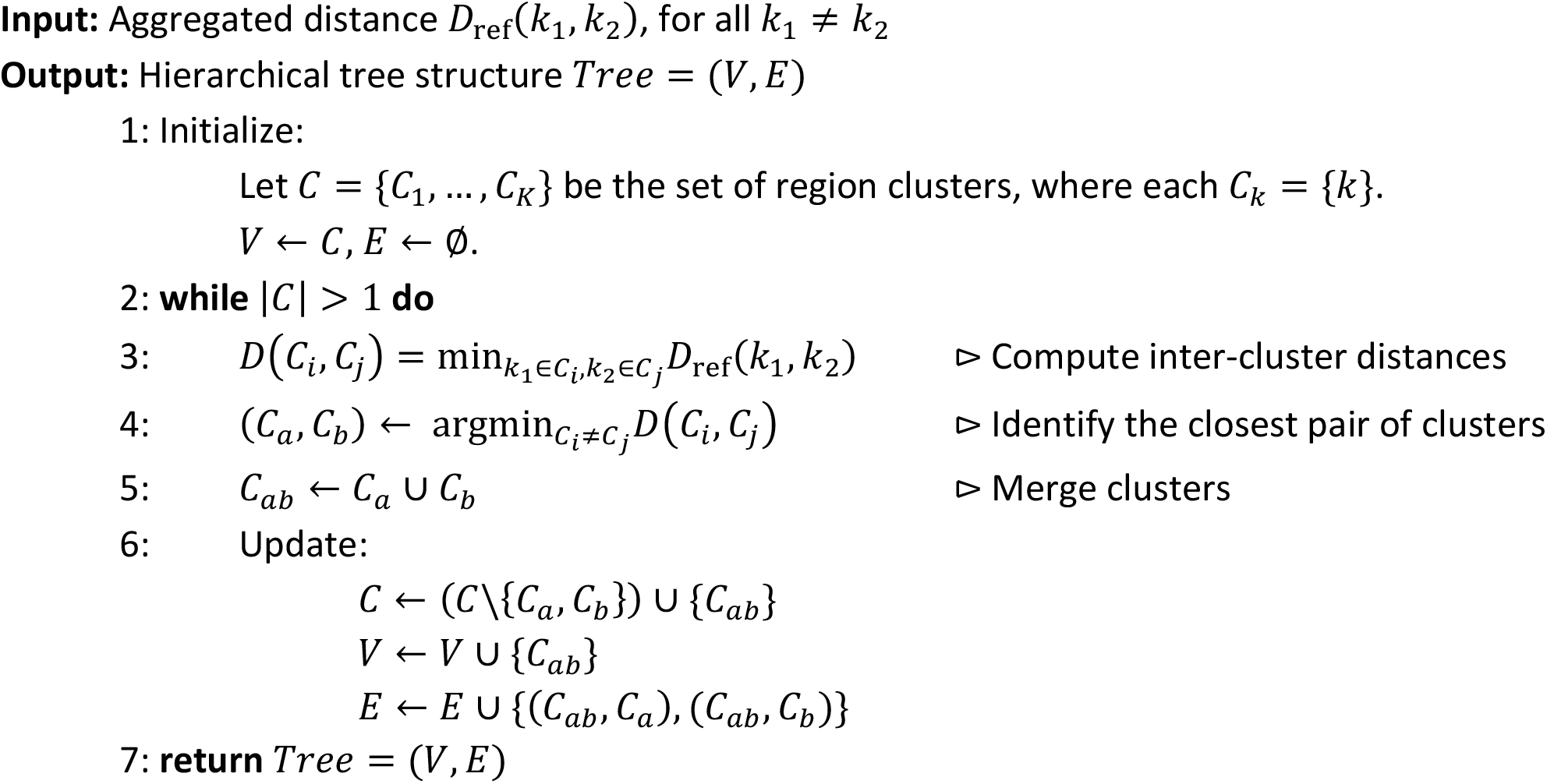

### Reference selection

For each query section *q* ∈ *S*_query_, HiCAT selects a subset of reference sections *S*_ref_(*q*) ⊂ *S*_ref_ that provide well-matched supervision. This query-specific reference selection enables the model to focus on the most relevant supervision and reduces noise from poorly matched references.

HiCAT quantifies reference-query compatibility using gene expression alone, as it is more consistently available across datasets than protein measurements and more stable across sections than H&E images, which are prone to staining artifacts. For a given query section *q*, we assess its similarity to each reference section *s* ∈ *S*_ref_ by comparing their gene expression distributions over the section-specific genes *f*_*s*_, using the two-sample Kolmogorov-Smirnov (KS) distance. Specifically, for each gene *g* ∈ *f*_*s*_, the KS distance is defined as:

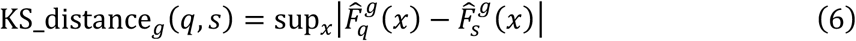

Where 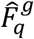 and 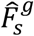denote the empirical cumulative distribution functions of gene *g* in sections *q* and *s*, respectively. We convert this distance into a similarity score KS_similarity_g_(*q, s*) = 1 − KS_distance_g_(*q, s*) ∈ [0,1], where larger values indicate greater similarity. The overall molecular similarity is defined as the average across all genes *g* ∈ *f*_*s*_.

In addition to molecular compatibility, references with a greater diversity of tissue region types are prioritized, as they can provide more comprehensive annotation coverage for the query. To account for this, HiCAT introduces a region coverage weight for each reference 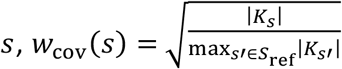, where |*K*_*s*_ | denotes the number of tissue regions in section *s*. The final similarity score between query *q* and reference *s* is defined as:

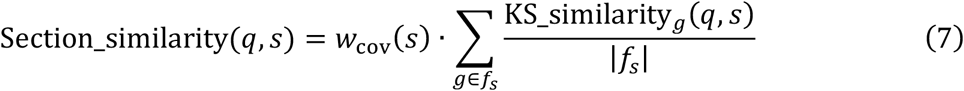

Next, HiCAT selects its reference set using relative thresholding:

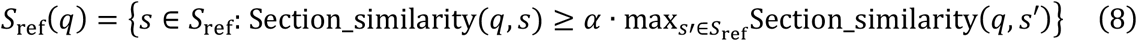

with *α* = 0.85 by default. This data-driven scheme accounts for tissue-specific variability, yielding larger reference sets in tissues with conserved transcriptomic profiles and more selective sets in heterogeneous tissues.

### Hierarchical label transfer

Multi-modal spatial omics provide complementary information, but their transferability across samples differs. Image features capture fine-grained structural patterns within individual sections but are more susceptible to artifacts and batch effects across samples. In contrast, molecular features are more biologically interpretable and transferable across samples but may not capture fine-grained structural patterns due to sequencing resolution and noise. Accordingly, HiCAT adopts a modality-aware integration strategy to ensure robust cross-sample label transfer.

#### Step 1: Modality selection

As modality informativeness varies across tissue types, HiCAT adopts a data-driven procedure to select modalities for clustering. Let *M* denotes the modalities included in the spatial dataset. For each modality *m* ∈ *M* and reference section *s* ∈ *S*_ref_, we consider the feature matrix 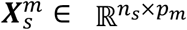. HiCAT reduces dimensionality to the top 50 principal components, followed by K-means clustering with |*K*_*s*_| clusters. Clustering performance is evaluated using the ARI by comparing the predicted assignment 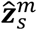 with ground-truth clusters ***z***_*s*_, derived from true labels ***y***_*s*_ by retaining partition structure while ignoring label identities. To quantify modality informativeness, we summarize performance across reference sections using the median ARIs. HiCAT exclude modalities whose performance below 0.85 times that of the best-performing modality, and denote the remaining modality set as *M*′. If all modalities exhibit low agreement with ground truth, HiCAT defaults to gene expression.

#### Step 2: Feature selection

Following the hierarchical tree *Tree* = (*V, E*), HiCAT performs feature selection separately for each bifurcating node.

For references and queries from the same study, features are selected within each reference section. For each non-leaf node 𝒱 ∈ *V* with two child nodes 𝒱_1_ and 𝒱_2_, and reference section 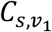 and 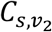denote the sets of tissue regions associated with 𝒱_1_ and 𝒱_2_, respectively. For each modality *m* ∈ *M*′, HiCAT identifies feature sets 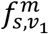and 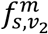that distinguish region clusters 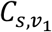and 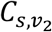using DE analysis with FDR-adjusted P value < 0.05, fold change > 1.15, top 10 features by fold change.

For reference and query from different studies, feature selection is performed jointly across all reference sections to identify shared signals. Feature-level DE statistics are first computed within each reference section and then aggregated across sections using the median. Features passing the filtering criterion are kept. This yields shared feature sets 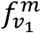 and 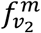that distinguish region clusters 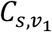 and 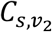.

These strategies reflect a scenario-adaptive design: independent selection preserves section-specific signals for within-study transfer, whereas joint selection captures shard features for robust cross-study/technology alignment.

#### Step 3: Clustering

HiCAT performs unsupervised clustering along the tree hierarchy, starting at the root and recursively partitioning at each bifurcating node until reaching the leaves. For each modality *m* ∈ *M*′ at a non-leaf node 𝒱 ∈ *V*, HiCAT considers two alternative embedding approaches: (1) a variance-preserving PCA embedding; and (2) a hierarchy-informed feature set defined as 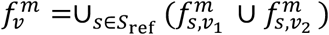, which aggregates discriminative signals across reference sections.

Next, HiCAT selects one embedding approach per modality and evaluates different combinations across modalities using the reference sections. For each combination, the selected modality-specific representations are concatenated into a joint feature space, and Louvain clustering is applied to obtain node-specific cluster assignments 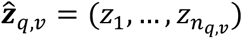,where *z*_*i*_ ∈ {1, …, *K*_*q,v*_} and 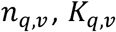denote the number of spots and clusters at node 𝒱 in query section *q*. HiCAT selects the combination with the highest ARI against the ground truth and applies the corresponding embedding strategy to the query samples.

#### Step 4: Anchor detection

HiCAT performs anchor detection in the molecular feature space, illustrated here using gene expression. To accommodate different scenarios, HiCAT uses two complementary strategies: a k-nearest-neighbor (kNN)-based framework for within-study label transfer, and a quantile-based framework for cross-study/technology transfer.

In the kNN-based approach, given a query section *q* and its selected references set *S*_ref_(*q*), HiCAT constructs a node-specific gene set 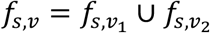 from each reference section *s* ∈ *S*_ref_(*q*) at every non-leaf node 𝒱∈ *V* with child nodes 𝒱_1_ and 𝒱_2_ . The corresponding feature matrix is defined as:

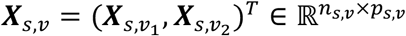

where *n*_*s*,𝒱_ denotes the number of spots assigned to node 𝒱 in reference *s, p*_*s*,𝒱_ = |*f*_*s*,𝒱_|, and 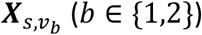 denotes the submatrix corresponding to spots assigned to child node 𝒱_b_. For the query section *q*, let 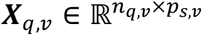 denote the feature matrix constructed using the same gene set.

Let 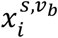 denote the expression vector of 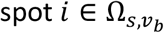 extracted from 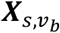, where 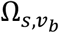 is the set of spots assigned to child node 𝒱_b_ in reference section *s*. For each reference spot *i*, HiCAT identifies its nearest neighbors among query spots assigned to node 𝒱 and define the spot-level anchor set as:

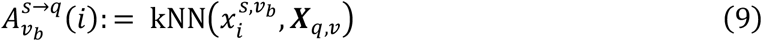

where kNN(∙) returns the set of *K*_*nn*_ = 5 nearest neighbors under Euclidean distance by default. Given a reference section *s*, anchors for child node 𝒱_b_ are aggregated across spots: 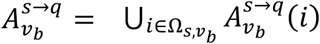. Anchor detection is performed independently by each reference section, and the final anchor set is aggregated across all selected reference sections: 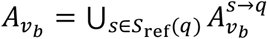.

For cross-study/technology transfer, batch effects and domain shifts can distort distance-based similarity in the kNN-based approach. To address this, HiCAT adopts a quantile-based framework that enables more robust anchor detection under domain shifts. Specifically, for each non-leaf node 𝒱∈ *V* with child nodes 𝒱_b_ (*b* = {1,2}), and for a given query section *q* with selected references *S*_ref_(*q*), HiCAT constructs node-specific gene expression matrices using the shared gene set across references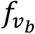 as:

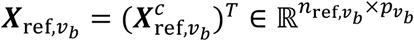

where 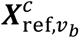 is the submatrix corresponding to spots in region 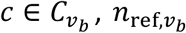 is the total number of spots associated with child node 𝒱_b_ across references, and 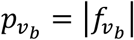.

For each gene 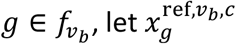 denote its expression vector in 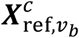. HiCAT first computes a region-specific lower expression bound as the 25^*th*^ percentile of expression values across reference spots within each region. The gene-level enrichment threshold is then defined as the median of these region-specific bounds across all regions:

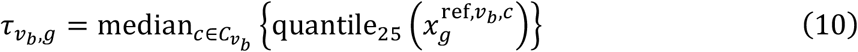

For the query section 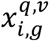 denote the expression vector of gene 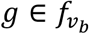 at spot 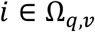.HiCAT identifies gene-level anchor evidence as 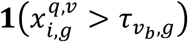. Aggregating across genes yields the gene enrichment score:

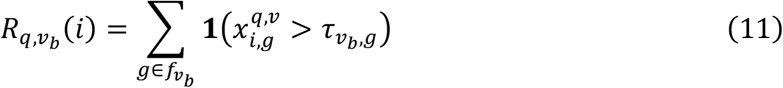

where 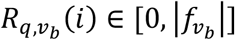 quantifies the strength of evidence supporting query spot *i* as a candidate anchor for child node 𝒱_b_. The final anchor set for child node 𝒱_b_ is defined as:

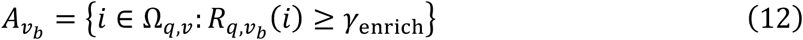

where *γ*_enrich_ is the enrichment score threshold and is determined in a data-driven manner as:

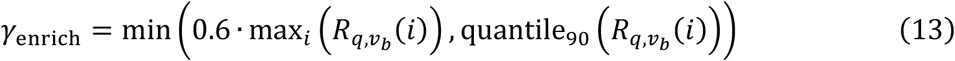

By requiring concordant enrichment across multiple discriminative genes, this approach improves robustness to batch effects and scale differences across datasets.

#### Step 5: Label assignment

Using the clustering results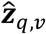 from **Step 3** and the anchor sets 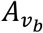 for each child node 𝒱_b_ (*b* = 1,2) from **Step 4**, HiCAT next computes the distribution of anchors across query clusters. Specifically, we define an anchor proportion matrix:

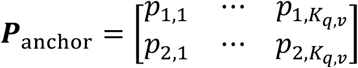

where *p*_*b,j*_ denotes the proportion of anchors from child node 𝒱_b_ assigned to query cluster *j*, with *j* ∈ {1, …, *K*_*q, 𝒱*_}. Each cluster is then assigned to the child node with the larger anchor proportion, resulting a binary partition at node 𝒱. This procedure is applied recursively along hierarchical tree until all region labels are determined. HiCAT also allows detection of novel clusters when anchor evidence is weak (e.g., maximum anchor proportion < 5%).

#### Step 6: Label enhancement

For low-resolution spatial data with H&E image, HiCAT includes an optional label enhancement step to improve boundary delineation. It leverages TESLA^48^ to integrate gene expression and histology images, generating pixel-level, resolution-enhanced ST data during preprocessing. The full pipeline is then applied to this enhanced data, enabling higher-resolution label assignment and finer-grained heterogeneity subtype characterization.

### Calculating heterogeneity score

To quantify cross-sample molecular heterogeneity within each annotated region, HiCAT defines a composite score considering the stability of region-specific markers and the divergence of gene expression profiles across samples.

For region *k*, let *S*_ref,*k*_ denote the set of reference sections containing the region. For *s* ∈ *S*_ref,*k*_,HiCAT identifies region-specific marker genes *f*_*s,k*_ using DE analysis. The similarity of marker gene sets between *s* and *s*′ is quantified by the Jaccard index: 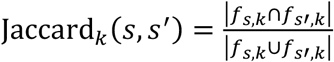 . The stability of marker gene set across samples is then quantified by the average of pairwise Jaccard index:

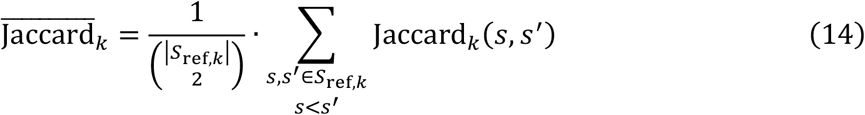

To obtain a gene-level heterogeneity measure while accounting for gene set size, we define:

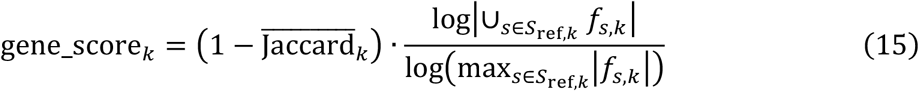

where a higher gene score indicates greater cross-sample heterogeneity in marker gene sets.

HiCAT next assesses heterogeneity considering the expression magnitude. Let 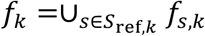 be the union gene set, and construct the region-specific expression matrix:

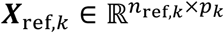

where *n*_*R*ef,*k*_ is the number of spots in region *k* across all reference sections and *p*_*k*_ = |*f*_*k*_|. Let 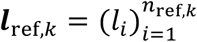, where ***l***_-_ ∈ *S*_ref,*k*_ denotes the sample identity of spot *i*. We quantify expression divergence across samples using the silhouette coefficient: silh_coef(***X***_*r*ef,*k*_,***l***_*r*ef,*k*_), where expression profiles are used as features and sample identities as labels. To evaluate whether the observed separation reflects true cross-sample heterogeneity, we generate a null distribution by permuting sample labels 200 times. The adjusted expression divergence score is defined as:

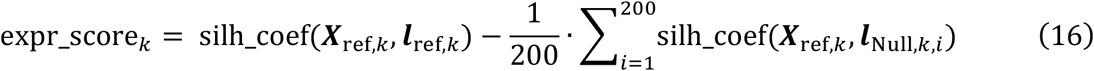

where ***l***_Null,*k,i*_ denotes the permuted labels in the *i*^th^ iteration.

The final heterogeneous score for region *k* is defined as:

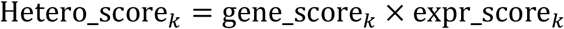

Scores are min-max normalized across regions to [0,1], with higher values indicate stronger cross-sample heterogeneity, reflecting both unstable region-specific marker gene sets and divergent expression profiles. Regions with normalized score > 0.5 are considered heterogeneous, with a permutation-derived P-values to assess deviation from random structure.

### Annotating heterogeneous subtypes

For heterogenous regions identified in the previous step, HiCAT further resolves shared molecular subtypes across samples. For each reference section *s* ∈ *S*_ref,*k*_, PCA is applied to the region-specific expression matrix, followed by Louvain clustering to identify within-sample subtype clusters. Subtype-specific marker genes are then identified via DE analysis within the section, and their union set defines a shared feature space capturing subtype-discriminative signals across samples.

To identify shared subtypes, all spots from region *k* across reference sections are represented in this feature space and clustered using K-Means with the number of clusters ranging from 2 to *N*_*k*_, where *N*_*k*_ denotes the total number of subtypes across samples. The number of shared subtypes is determined by maximizing the average silhouette coefficient. The resulting clusters define shared molecular subtypes across samples. For each shared subtype, HiCAT pool spots across sections and perform DE analysis to identify subtype-specific marker genes, enabling downstream functional characterization.

These subtype assignments are then mapped back to spatial coordinates to yield refined, high-resolution annotations that capture intra-region molecular heterogeneity beyond pathologist labels. The identified subtype signatures are further used to propagate subtype labels to query samples by the quantile-based anchor detection framework, enabling consistent subtype identification across cohort.

### Benchmark analyses

For supervised benchmarking, we compared HiCAT with state-of-the-arts supervised methods (Seurat, SpaDo, SCGP, STELLAR), evaluated using accuracy and F1 score. For unsupervised benchmarking, we evaluated against six unsupervised methods (IRIS, BASS, BayesSpace, STAGATE, Seurat, SCGP) using ARI. Implementation details, including preprocessing, parameter settings, post-processing, and reference usage for supervised methods, are summarized in **Supplementary Table 5 and 6**.

## Data availability

We analyzed six spatial transcriptomics data and one spatial omics data, together with three matched scRNA-seq reference datasets for benchmarking. Publicly available data were acquired from the following websites or accession numbers: (1) human HER2-positive breast tumor spatial transcriptomics data (https://github.com/almaan/her2st); (2) human breast tumor 10x Visium data (https://www.10xgenomics.com/datasets/human-breast-cancer-visium-fresh-frozen-whole-transcriptome-1-standard); (3) human breast tumor spatial transcriptomics data (https://data.mendeley.com/datasets/29ntw7sh4r/5); (4) human tonsil 10x Visium gene and protein expression data (sample 1: https://www.10xgenomics.com/datasets/visium-cytassist-gene-and-protein-expression-library-of-human-tonsil-with-add-on-antibodies-h-e-6-5-mm-ffpe-2-standard; sample 2: https://www.10xgenomics.com/datasets/gene-protein-expression-library-of-human-tonsil-cytassist-ffpe-2-standard); (5) human tonsil 10x Visium HD data (https://www.10xgenomics.com/datasets/visium-hd-cytassist-gene-expression-human-tonsil-fresh-frozen); (6) mouse brain 10x Visium data (https://www.10xgenomics.com/cn/datasets/multiomic-integration-neuroscience-application-note-visium-for-ffpe-plus-immunofluorescence-alzheimers-disease-mouse-model-brain-coronal-sections-from-one-hemisphere-over-a-time-course-1-standard); (7) human prostate tumor 10x Visium data (https://data.mendeley.com/datasets/svw96g68dv/1); (8) human breast tumor scRNA-seq reference data (GEO accession: GSE176078); (9) human tonsil scRNA-seq reference data (https://zenodo.org/records/10373041); (10) mouse brain scRNA-seq reference data (http://mousebrain.org/adolescent/).

Details of the datasets analyzed in this paper were summarized in **Supplementary Table 7**.

## Software availability

An open-source implementation of the HiCAT algorithm can be downloaded from GitHub: https://github.com/jinghuang-stats/HiCAT

## Life sciences reporting summary

Further information on experimental design is available in the Life Sciences Reporting Summary.

