## Supplementary information for "From unsupervised clustering to atlas-guided annotation in cohort-scale spatial omics with HiCAT"

**Supplementary Table 1. Training references for supervised method in the HER2-postive breast cancer dataset**

| **Method** | **Reference integration** | **Available references** | **Query section** | **References used for benchmarking** |
|---|---|---|---|---|
| HiCAT | Multi-reference | Subsets of the reference pool {E1, G2, H1} | A1 | G2 |
| | | | B1 | G2 + H1 |
| | | | C1 | E1 + G2 + H1 |
| | | | D1 | E1 + G2 + H1 |
| | | | F1 | E1 + G2 + H1 |
| Seurat | Multi-reference | {E1}, {G2}, {H1}, {E1 + G2 + E1} | A1 | G2 |
| | | | B1 | E1 + G2 + H1 |
| | | | C1 | E1 + G2 + H1 |
| | | | D1 | G2 |
| | | | F1 | G2 |
| SpaDo | Single-reference | {E1}, {G2}, {H1} | A1 | G2 |
| | | | B1 | H1 |
| | | | C1 | G2 |
| | | | D1 | H1 |
| | | | F1 | G2 |
| SCGP | Multi-reference | {E1}, {G2}, {H1}, {E1 + G2 + H1} | A1 | H1 |
| | | | B1 | H1 |
| | | | C1 | G2 |
| | | | D1 | G2 |
| | | | F1 | H1 |
| STELLAR | Single-reference | {E1}, {G2}, {H1} | A1 | E1 |
| | | | B1 | H1 |
| | | | C1 | E1 |
| | | | D1 | E1 |
| | | | F1 | E1 |

All methods were provided with the same candidate reference pool {E1, G2, H1}. “Available references” denotes the set of reference inputs each method can accept. HiCAT incorporates an automatic pipeline to select suitable references from the pool for each query section.

To ensure a fair comparison, although the benchmarking methods do not include an explicit reference selection procedure, they were evaluated across all feasible reference configurations supported by each method. This includes each individual reference section and, for methods capable of integrating multiple references, the full reference pool as a combined input. The best-performing references for each method were reported and used for benchmarking evaluation.

**Supplementary Table 2. Training references for supervised method in the 10x Visium breast cancer dataset**

| Method | Reference integration | Available references | Query section | References used for benchmarking |
|---|---|---|---|---|
| HiCAT | Multi-reference | Subsets of the reference pool {A1, B1, C1, D1, E1, F1, G2, H1} | 10x BC | B1 + G2 + H1 |
| Seurat | Multi-reference | {A1}, {B1}, {C1}, {D1}, {E1}, {G2}, {H1}, {A1 + B1 + C1 + D1 + E1 + G2 + H1} | 10x BC | A1 + B1 + C1 + D1 + E1 + F1 + G2 + H1 |
| SpaDo | Single-reference | {A1}, {B1}, {C1}, {D1}, {E1}, {G2}, {H1} | 10x BC | F1 |
| SCGP | Multi-reference | {A1}, {B1}, {C1}, {D1}, {E1}, {G2}, {H1}, {A1 + B1 + C1 + D1 + E1 + G2 + H1} | 10x BC | G2 |
| STELLAR | Single-reference | {A1}, {B1}, {C1}, {D1}, {E1}, {G2}, {H1} | 10x BC | C1 |

All methods were provided with the same candidate reference pool {A1, B1, C1, D1, E1, F1, G2, H1}. The supervised benchmarking setup follows that described in **Supplementary Table 1**.

**Supplementary Table 3. Training references for supervised method in the breast cancer ST cohort**

| Method | Reference integration | Available references | Query breast cancer subtype | Query section | Selected references |
|---|---|---|---|---|---|
| HiCAT | Multi-reference | Subsets of the reference pool {A1, B1, C1, D1, E1, F1, G2, H1} | TNBC | BC23288_D2 | B1 + G2 + H1 |
| | | | TNBC | BC23377_C2 | A1 + B1 + G2 + H1 |
| | | | TNBC | BC23803_D2 | A1 + C1 + D1 + E1 + G2 + H1 |
| | | | TNBC | BC23209_D1 | A1 + G2 + H1 |
| | | | HER2 luminal | BC23450_D2 | G2 + H1 |
| | | | HER2 luminal | BC23901_C2 | B1 + G2 + H1 |
| | | | HER2 luminal | BC24220_E1 | A1 + B1 + G2 + H1 |
| | | | HER2 luminal | BC23287_D1 | B1 + H1 |
| | | | HER2 luminal | BC23944_D2 | B1 + G2 + H1 |
| | | | LuminalB | BC23508_E2 | B1 + G2 + H1 |
| | | | LuminalB | BC23277_D2 | B1 + G2 + H1 |
| | | | LuminalB | BC23506_C2 | A1 + C1 + D1 + E1 + G2 + H1 |
| | | | LuminalB | BC23270_D2 | B1 + G2 + H1 |
| | | | LuminalB | BC23895_C1 | A1 + B1 + C1 + D1 + E1 + G2 + H1 |
| | | | LuminalA | BC23272_E1 | B1 + G2 + H1 |
| | | | LuminalA | BC23269_C1 | B1 + G2 + H1 |
| | | | LuminalA | BC24223_D2 | B1 + G2 + H1 |
| | | | LuminalA | BC23268_C2 | B1 + G2 + H1 |

**Supplementary Table 4. Training references used for supervised method in the mouse brain dataset**

| Method | Query section | References used for benchmarking |
|---|---|---|
| HiCAT | WT 3m rep2 | WT 3m rep1 + WT 6m rep1 |
| | WT 6m rep2 | WT 3m rep1 + WT 6m rep1 + WT 13m rep1 |
| | WT 13m rep2 | WT 13m rep1 |
| | AD 3m rep2 | AD 3m rep1 |
| | AD 6m rep2 | AD 3m rep1 + AD 6m rep1 + AD 18m rep1 |
| | AD 18m rep2 | AD 6m rep1 + AD 18m rep1 |
| Seurat | WT 3m rep2 | WT 3m rep1 |
| | WT 6m rep2 | WT 6m rep1 |
| | WT 13m rep2 | WT 13m rep1 |
| | AD 3m rep2 | AD 3m rep1 |
| | AD 6m rep2 | AD 6m rep1 |
| | AD 18m rep2 | AD 18m rep1 |
| SpaDo | WT 3m rep2 | WT 3m rep1 |
| | WT 6m rep2 | WT 6m rep1 |
| | WT 13m rep2 | WT 13m rep1 |
| | AD 3m rep2 | AD 3m rep1 |
| | AD 6m rep2 | AD 6m rep1 |
| | AD 18m rep2 | AD 18m rep1 |
| SCGP | WT 3m rep2 | WT 3m rep1 |
| | WT 6m rep2 | WT 6m rep1 |
| | WT 13m rep2 | WT 13m rep1 |
| | AD 3m rep2 | AD 3m rep1 |
| | AD 6m rep2 | AD 6m rep1 |
| | AD 18m rep2 | AD 18m rep1 |
| STELLAR | WT 3m rep2 | WT 3m rep1 |
| | WT 6m rep2 | WT 6m rep1 |
| | WT 13m rep2 | WT 13m rep1 |
| | AD 3m rep2 | AD 3m rep1 |
| | AD 6m rep2 | AD 6m rep1 |
| | AD 18m rep2 | AD 18m rep1 |

WT denotes wild type and AD denotes Alzheimer's disease; rep1/rep2 indicate biological replicate1/replicate2. In the mouse brain dataset, two biological replicates are available for each time point and condition. For benchmark methods, the paired biological replicate was used as the reference for each query sample, consistent with their standard pipeline design. For HiCAT, query samples were provided with condition-matched reference pools (WT: {WT 3m rep1, WT 6m rep1, WT 13m rep1}; AD: {AD 3m rep1, AD 6m rep1, AD 18m rep1}), allowing automatic selection of suitable references within each condition.

**Supplementary Table 5. Supervised benchmark methods implementation details**

<table>
<tr><th>Method</th><th>Preprocessing</th><th>Parameter settings</th></tr>
<tr><td>Seurat[1]</td><td>Follow default Seurat preprocessing steps:<br>1. Log-normalization: counts normalized to 10,000 per spot, followed by $\log(1 + normalized\ counts)$ transformation;<br>2. Variable feature selection: top 2,000 highly variable genes;<br>3. Scaling: data scaled with a maximum value at 10;<br>4. Dimensionality reduction: top 50 principal components (PCs) retained.</td><td>Default settings</td></tr>
<tr><td>SpaDo[2]</td><td>Follow default SpaDo preprocessing steps:<br>For spot-level data, SpaDo requires scRNA-seq data from the same tissue type as references to obtain cell type deconvolution using Cell2Location:<br>1. Reference signature estimation: infer cell type-specific gene expression profiles from scRNA-seq data;<br>2. Spatial mapping: estimate the proportion of each cell type within each spatial spot.<br><br>For cell-level data:<br>1. Log-normalization: counts normalized to 10,000 per cell, followed by $\log(1 + normalized\ counts)$ transformation.</td><td>Default settings</td></tr>
<tr><td>SCGP[3]</td><td rowspan="2">SCGP and STELLAR do not contain a default preprocessing steps; therefore, we adopt commonly used preprocessing steps for spatial transcriptomics data:<br>1. Gene filtering: remove genes that have nonzero expression lower than 5%;<br>2. Log-normalization: counts normalized to 10,000 per cell, followed by $\log(1 + normalized\ counts)$ transformation.<br>3. Dimensionality reduction: top 40 principal components (PCs) retained.</td><td>The resolution parameter $rp$ controls the number of resulting clusters. By default, $rp = 0.0001$; however, this setting can yield too many clusters in some datasets. To ensure a fair and meaningful comparison, we manually adjust rp within the range of 0.01-0.00005 to obtain a number of clusters comparable to the ground-truth region categories across datasets. This tuning improves SCGP performances in the benchmarking.</td></tr>
<tr><td>STELLAR[4]</td><td>Default settings</td></tr>
</table>

**Supplementary Table 6. Unsupervised benchmark methods implementation details**

| Method | Preprocessing | Parameter settings |
|---|---|---|
| IRIS[5] | Follow default IRIS preprocessing steps:<br>1. Gene filtering: filter out genes expressed in fewer than 5 spots;<br>2. Spots filtering: filter out spots with fewer than 100 detected genes. | To control the number of resulting spatial clusters, set $numCluster$ to the number of ground truth labels. |
| SCGP[3] | SCGP do not contain a default preprocessing steps; therefore, we adopt commonly used preprocessing steps for spatial transcriptomics data:<br>1. Gene filtering: remove genes that have nonzero expression lower than 5%;<br>2. Log-normalization: counts normalized to 10,000 per cell, followed by $\log(1 + normalized\ counts)$ transformation.<br>3. Dimensionality reduction: top 40 PCs retained. | By default, $rp = 0.0001$. We adjust $rp$ within the range of 0.01-0.00005 to obtain a number of clusters comparable to the ground-truth region categories across datasets. |
| BayesSpace[6] | Follow default BayesSpace preprocessing steps:<br>1. Log-normalization: counts normalized by library size (per spot) followed by $\log2(normalized\ counts)$;<br>2. Variable feature selection: top 2,000 highly variable genes;<br>3. Dimensionality reduction: top 15 PCs retained. | To control the number of resulting spatial clusters, set $q$ to the number of ground truth labels. |
| BASS[7] | Follow default BASS preprocessing steps:<br>1. Log-normalization: counts normalized by library size (per spot) followed by $\log2(normalized\ counts)$;<br>2. Variable feature selection: top 3,000 highly variable genes;<br>3. Dimensionality reduction: top 20 PCs retained. | To control the number of resulting spatial clusters, set $R$ to the number of ground truth labels. |
| STAGATE[8] | Follow default STAGATE preprocessing steps:<br>1. Log-normalization: counts normalized to 10,000 per spot, followed by $\log(1 + normalized\ counts)$ transformation;<br>2. Variable feature selection: top 3,000 highly variable genes; | To control the number of resulting spatial clusters, set $num\_cluster$ to the number of ground truth labels. |
| Seurat[1] | Follow default Seurat preprocessing steps:<br>1. Log-normalization: counts normalized to 10,000 per spot, followed by $\log(1 + normalized\ counts)$ transformation;<br>2. Variable feature selection: top 2,000 highly variable genes;<br>3. Scaling: data scaled with a maximum value at 10;<br>4. Dimensionality reduction: top 50 PCs retained. | By default, $resolution = 0.8$. We adjust $resolution$ within the range of 0.1-1.2 to obtain a number of clusters comparable to the ground-truth region categories across datasets. |

**Supplementary Table 7. Summary of analyzed datasets**

| Tissue | Data source | Sample ID | Number of spots/cells | Number of genes/proteins | Protocol |
|---|---|---|---|---|---|
| Human HER2-positive breast tumor | Andersson *et al.*[9] (https://github.com/almaan/her2st) | A1 | 346 | 15,045 genes | Spatial Transcriptomics |
| | | B1 | 295 | 15,109 genes | |
| | | C1 | 176 | 15,557 genes | |
| | | D1 | 306 | 15,661 genes | |
| | | E1 | 587 | 15,701 genes | |
| | | F1 | 691 | 14,861 genes | |
| | | G2 | 467 | 15,258 genes | |
| | | H1 | 613 | 15,029 genes | |
| Human breast tumor | 10x Genomics (https://www.10xgenomics.com/datasets/human-breast-cancer-visium-fresh-frozen-whole-transcriptome-1-standard) | NA | 4898 | 36,601 genes | 10x Visium |
| Human breast tumor | Stahl *et al.*[10] (https://data.mendeley.com/datasets/29ntw7sh4r/5) | BC23450_D2 | 316 | 19,391 genes | Spatial Transcriptomics |
| | | BC23901_C2 | 292 | 18,082 genes | |
| | | BC24220_E1 | 428 | 17,273 genes | |
| | | BC23287_D1 | 357 | 16,826 genes | |
| | | BC23944_D2 | 414 | 18,201 genes | |
| | | BC23272_E1 | 578 | 17,593 genes | |
| | | BC23269_C1 | 434 | 17,779 genes | |
| | | BC24223_D2 | 368 | 17,428 genes | |
| | | BC23268_C2 | 498 | 17,467 genes | |
| | | BC23508_E2 | 504 | 18,535 genes | |
| | | BC23277_D2 | 506 | 18,191 genes | |
| | | BC23506_C2 | 502 | 19,088 genes | |
| | | BC23270_D2 | 284 | 17,940 genes | |
| | | BC23895_C1 | 498 | 17,686 genes | |
| | | BC23288_D2 | 464 | 17,387 genes | |
| | | BC23377_C2 | 687 | 18,306 genes | |
| | | BC23803_D2 | 402 | 19,729 genes | |
| | | BC23209_D1 | 331 | 18,298 genes | |

| | | | | | |
|---|---|---|---|---|---|
| Human tonsil | 10x Genomics (https://www.10xgenomics.com/datasets/visium-cytassist-gene-and-protein-expression-library-of-human-tonsil-with-add-on-antibodies-h-e-6-5-mm-ffpe-2-standard) | sample 1 | 4,908 | 18,085 genes<br>41 proteins | 10x Visium Omics |
| | 10x Genomics (https://www.10xgenomics.com/datasets/gene-protein-expression-library-of-human-tonsil-cytassist-ffpe-2-standard) | sample 2 | 4,194 | 18,085 genes<br>35 proteins | |
| Human tonsil | 10x Genomics (https://www.10xgenomics.com/datasets/visium-hd-cytassist-gene-expression-human-tonsil-fresh-frozen) | NA | 175,448 | 18,085 genes | 10x Visium HD |
| Mouse brain | 10x Genomics (https://www.10xgenomics.com/cn/datasets/multiomic-integration-neuroscience-application-note-visium-for-ffpe-plus-immunofluorescence-alzheimers-disease-mouse-model-brain-coronal-sections-from-one-hemisphere-over-a-time-course-1-standard) | WT 3m rep1 | 2,960 | 19,465 genes | 10x Visium |
| | | WT 3m rep2 | 2,884 | 19,465 genes | |
| | | WT 6m rep1 | 3,361 | 19,465 genes | |
| | | WT 6m rep2 | 2,827 | 19,465 genes | |
| | | WT 18m rep1 | 3,621 | 19,465 genes | |
| | | WT 18m rep2 | 3,166 | 19,465 genes | |
| | | AD 3m rep1 | 2,955 | 19,465 genes | |
| | | AD 3m rep2 | 2,084 | 19,465 genes | |
| | | AD 6m rep1 | 3,296 | 19,465 genes | |
| | | AD 6m rep2 | 3,063 | 19,465 genes | |
| | | AD 18m rep1 | 3,427 | 19,465 genes | |
| | | AD 18m rep2 | 3,157 | 19,465 genes | |
| Human prostate tumor | Erickson *et al*.[11] (https://data.mendeley.com/datasets/svw96g68dv/1) | H2_1 | 3,092 | 34,562 genes | 10x Visium |
| | | H1_4 | 4,075 | 34,562 genes | |
| | | H1_5 | 3,837 | 34,562 genes | |
| Human breast tumor | Wu *et al*.[12] (https://www.ncbi.nlm.nih.gov/geo/query/acc.cgi?acc=GSE176078) | NA | 100,064 | 29,733 genes | scRNA-seq |
| Human tonsil | Massoni-Badosa *et al*.[13] (https://zenodo.org/records/10373041) | NA | 263,286 | 37,378 genes | scRNA-seq |

| | | | | | |
|---|---|---|---|---|---|
| Mouse brain | Zeisel *et al*.[14] (http://mousebrain.org/adolescent/) | NA | 32,068 | 27,933 genes | scRNA-seq |

**Supplementary Note 1: Region-specific detection performance across supervised methods in a HER2-positive breast cancer dataset**

In addition to overall annotation accuracy, we evaluated region-specific detection performance across supervised methods to assess how effectively each method identified individual tissue section. This comparison is important because overall accuracy can be dominated by large or prevalent regions, potentially obscuring poorer performance in smaller or more intricate regions.

For evaluation, we focused on region-specific precision and F1 score. Precision quantifies detection specificity by measuring the proportion of spots predicted as a given region that are correctly assigned. This metric is particularly important for spatial annotation, where misassignment of predicted regions into neighboring tissue can blur anatomical boundaries and affect downstream interpretations. F1 score provides a complementary measure that balances precision and recall.

As shown in **Supplementary Fig. 1a** and **1b**, HiCAT achieves the highest region-specific precision and F1 score across individual regions among all supervised methods. Compared with the second-best method, Seurat, HiCAT improved region-specific detection precision by 23-238% across regions, with a mean improvement of 105%. In contrast, Seurat showed weaker performance in detecting connective tissue, whereas SpaDo and SCGP struggled with small-region detection and with distinguishing *in situ* cancer from invasive cancer. STELLAR performed poorly across most regions, with only moderate performance in detecting invasive cancer.

Together, these results demonstrate that HiCAT provides more accurate region-specific annotation across diverse tissue structures, enabling reliable detection of major regions and finer delineation of smaller, biologically meaningful structures.

**Supplementary Fig. 1 |** Bar plots comparing region-specific **a.** precision and **b.** F1 score across supervised methods in HER2-positive breast cancer dataset, with the bar height indicating the mean metric value across query sections.

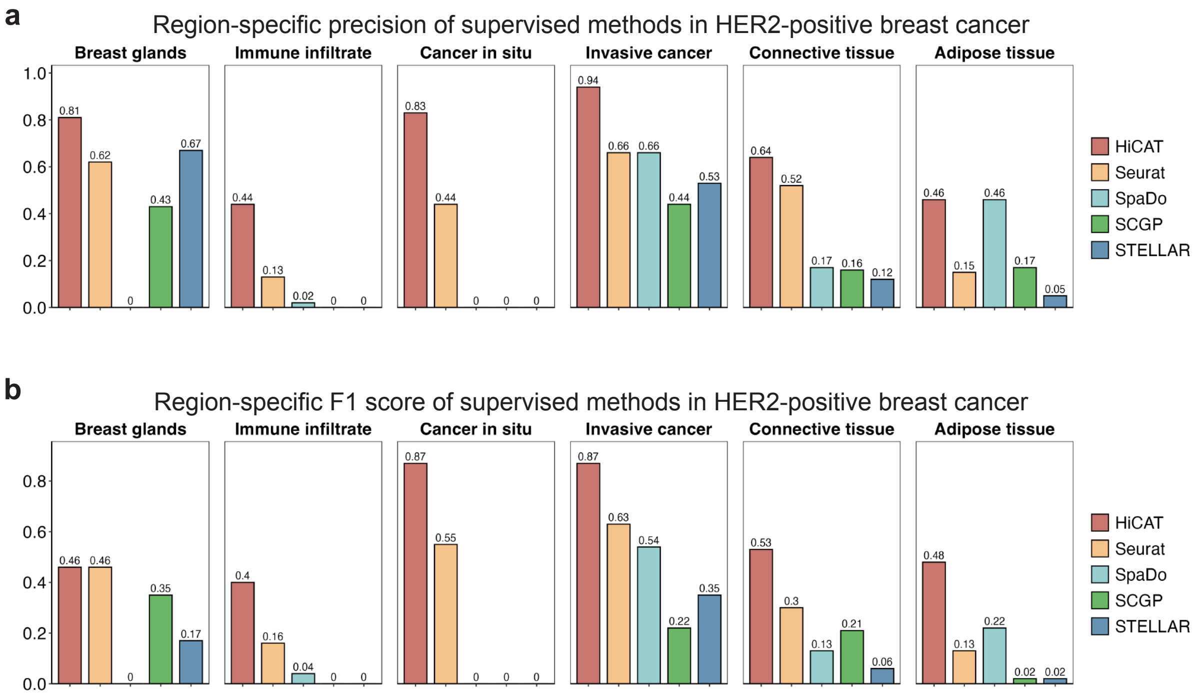

**Supplementary Note 2: Unsupervised benchmarking on a HER2-positive breast cancer dataset**

In addition to benchmarking against supervised methods, we compared HiCAT with six state-of-the-art unsupervised spatial domain detection methods, including IRIS, SCGP (unsupervised), Seurat (unsupervised), BayesSpace, BASS, and STAGATE. Method-specific parameter settings are detailed in **Supplementary Table 6**, and performance was evaluated using ARI.

**Supplementary Fig. 2a** shows that domain detection results across methods. HiCAT captures both broad region organization, such as the separation between tumor and normal regions, and finer anatomical structures, including small breast glands (cluster 6) and immune infiltrates (cluster 3). In contrast, competing unsupervised methods exhibited distinct domain detection patterns. Seurat and SCGP produced sparse and less spatially coherent domains, while BASS captured major region-level separation but introduced additional subclusters within broad tissue regions. BayesSpace, IRIS, and STAGATE tended to generate irregular clusters that deviated from annotated tissue boundaries and did not consistently reflect biologically meaningful region clusters.

As shown in **Supplementary Fig. 2b**, HiCAT achieved the highest average performance, with a mean ARI of 0.44, corresponding to 29%-340% improvement over the comparison methods. Although HiCAT had slightly lower ARI than the best competing method in sections B1 and C1, visual inspection of **Supplementary Fig. 2a** showed that, in B1, HiCAT detected small, spatially discrete breast gland structures that were not captured by BASS, reducing agreement with the broader region-level labels. In C1, HiCAT further resolved subregions within the annotated tumor region, which reduced ARI.

Together, these results show that HiCAT outperforms existing unsupervised spatial domain detection methods in recovering biologically meaningful tissue structures.

**Supplementary Fig. 2 | a.** Spatial domain detection results across HER2-positive breast cancer query sections. **b.** Bar plots comparing spatial domain detection performance across methods, evaluated by ARI. Bar heights indicate the mean ARI across query sections, and error bars represent standard errors.

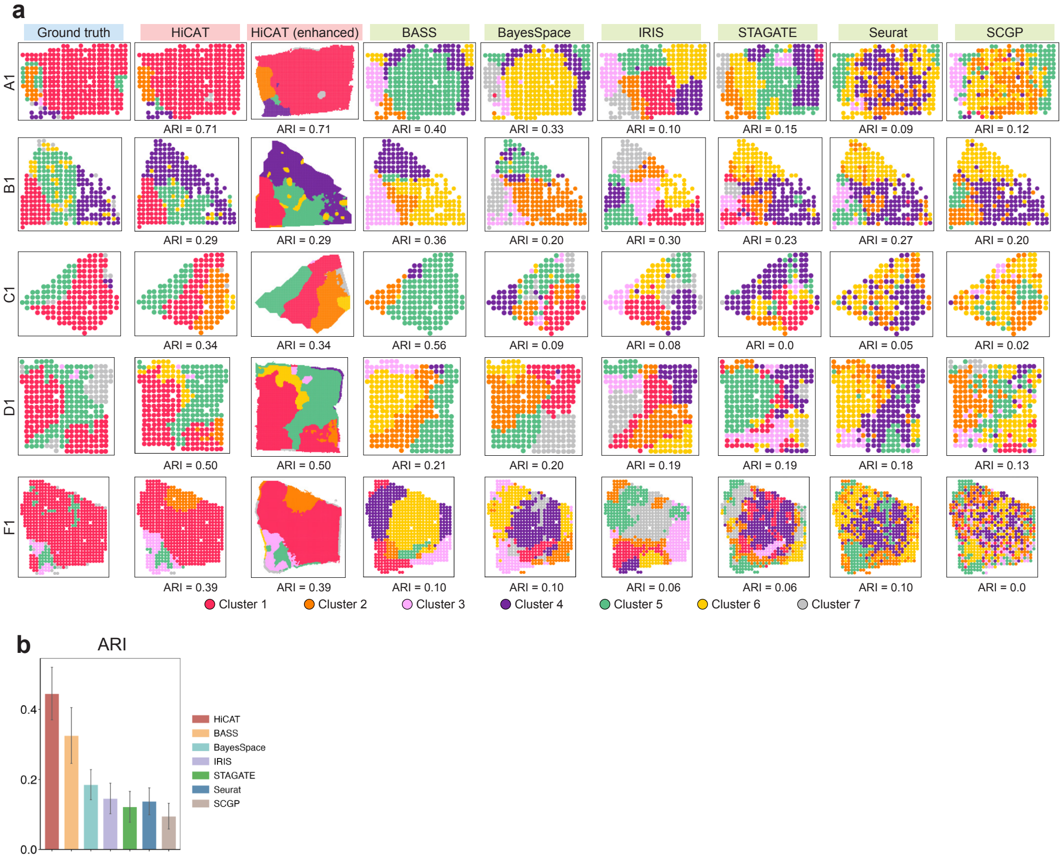

**Supplementary Fig. 3 |** Bar plots comparing region-specific **a.** precision and **b.** F1 score across supervised methods in breast cancer Visium dataset, with the bar height indicating the mean metric value across query sections.

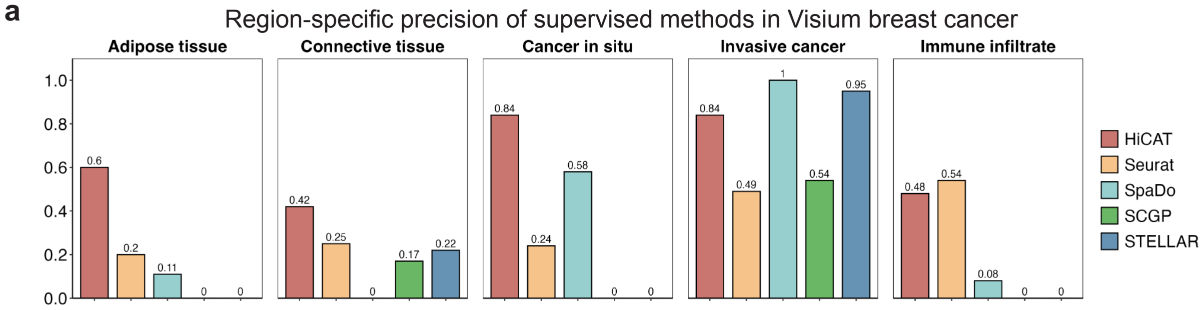


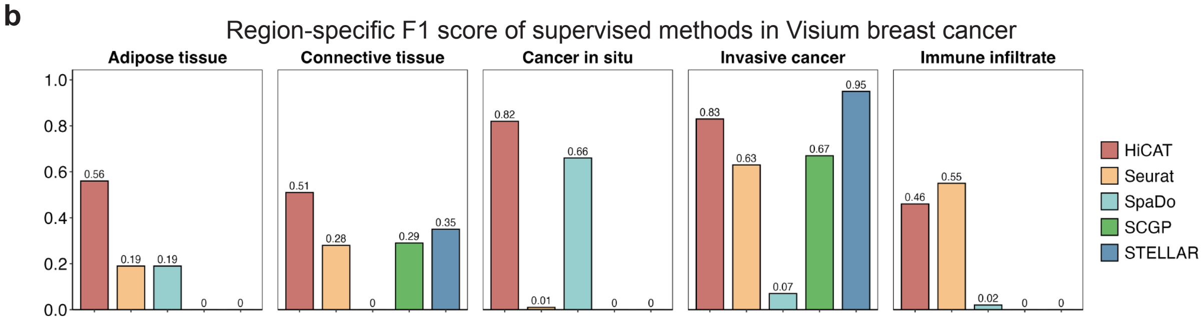

**Supplementary Note 3: Unsupervised benchmarking on a breast cancer Visium dataset**

We also benchmarked HiCAT against six unsupervised methods on the breast cancer Visium dataset. **Supplementary Fig. 2a** shows the spatial domain detection results across methods. HiCAT identified major tumor regions, distinguishing *in situ* (Cluster 2) from invasive (Cluster 1) tumor subtypes, and detected fine-grained small structures. In contrast, competing methods failed to identify small adipose tissue (Cluster 3) and connective tissue (Cluster 5), tended to split spatially discrete *in situ* cancer regions into separate clusters, identified additional subclusters within invasive cancer, and showed difficulty detecting immune infiltrate region (Cluster 4). As shown in **Supplementary Fig. 2b**, HiCAT achieved the highest domain detection performance, with an ARI of 0.51, corresponding to an 11%-131% improvement over the comparison methods.

**Supplementary Fig. 4 | a.** Spatial domain detection results for the breast cancer Visium section. **b.** Bar plots comparing spatial domain detection performance across methods, evaluated by ARI.

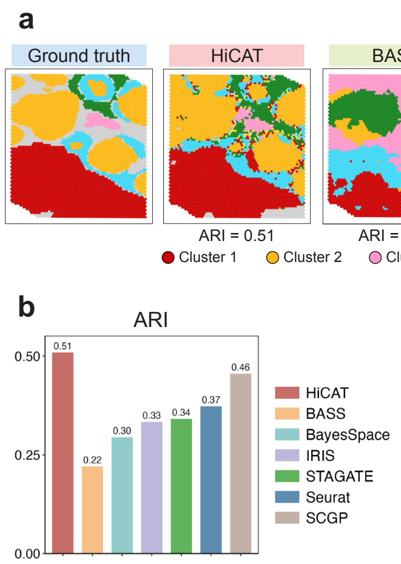

**Supplementary Note 4: Annotation of tumor and immune heterogeneity in HER2-positive breast cancer query samples**

Using tumor and immune subtypes identified in HER2-positive reference samples, together with subtype-specific markers shared across samples, we annotated tumor and immune subtypes in the HER2-positive query samples. HiCAT resolved fine-grained, enhanced-resolution subtypes within the identified tumor and immune regions through label transfer (**Supplementary Fig. 5a**). Immune infiltrate regions in A1 and D1 comprised less than 1% of spots and were therefore excluded from immune subtype annotation.

The hot tumor subtype showed enrichment of antigen-presentation and immune-activation programs, including MHC-II-associated genes (*HLA-DRA*, *HLA-DRB1*, *HLA-DQA1*, and *HLA-DPB1*) and interferon- or inflammatory response-related genes (*BST2*, *SOCS1*, *PSMBB*, *WARS*, and *HLA-B*) (**Supplementary Fig. 5b**). These patterns were consistent with pathway enrichment observed in the HER2-positive reference samples.

**Supplementary Fig. 5 | a.** Enhanced-resolution annotation plots showing the spatial distribution of fine-grained tumor and immune subtypes across HER2-positive breast cancer query samples. **b.** Gene expression enrichment heatmaps showing shared molecular programs distinguishing hot tumor from normal tumor regions across query sections.

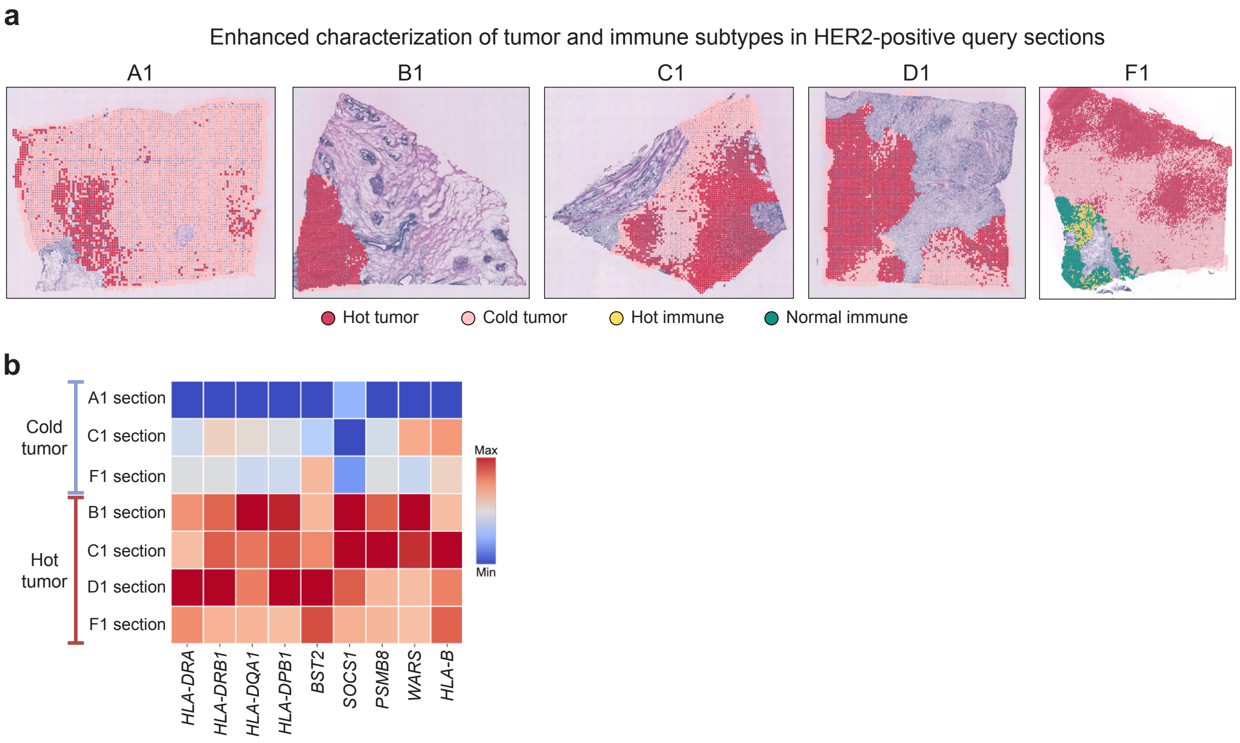

**Supplementary Fig. 6 | a.** Tissue region annotation in breast cancer ST cohort. HiCAT provides both spot-level and enhanced-resolution delineation of tissue regions across breast cancer clinical subtypes.

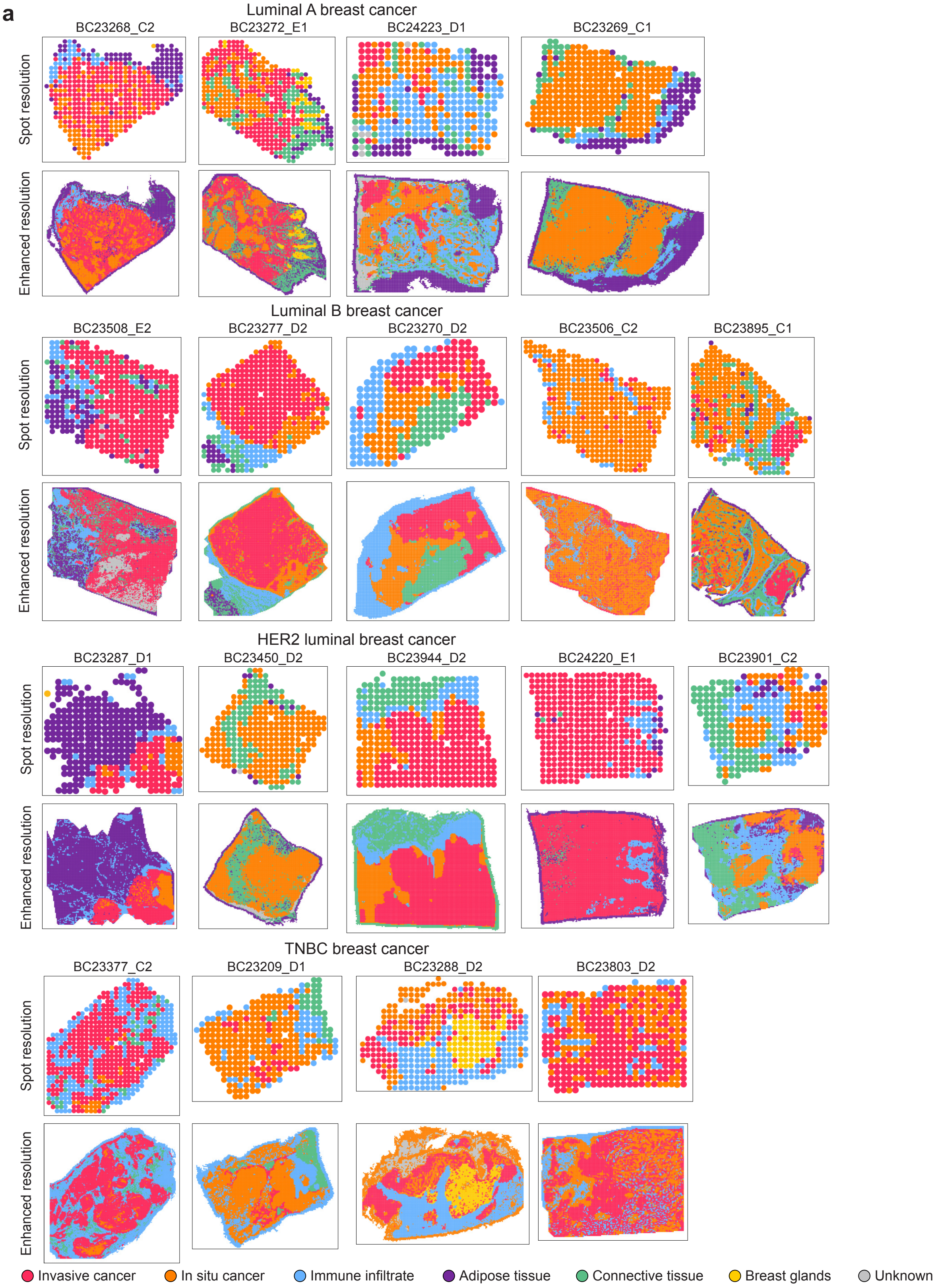

**Supplementary Fig. 7 |** Gene expression enrichment heatmaps illustrating shared molecular programs across breast cancer ST cohort. **a.** immune-inflamed, “hot” tumor state. **b.** activated, “hot” immune niche.

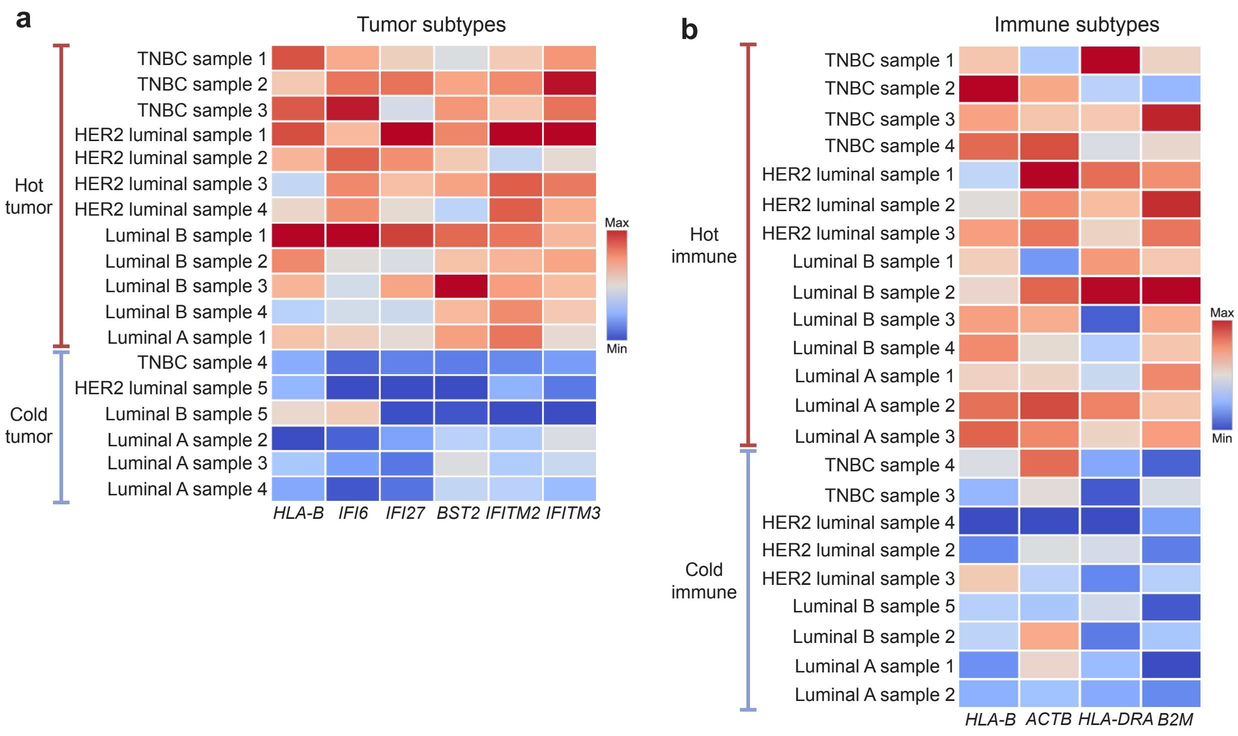

**Supplementary Fig. 8 | a.** Bar plots showing the expression levels of proliferation-related genes within tumor regions across breast cancer clinical subtypes. Bar heights indicate the mean expression across samples within each clinical subtype, and error bars represent standard errors.

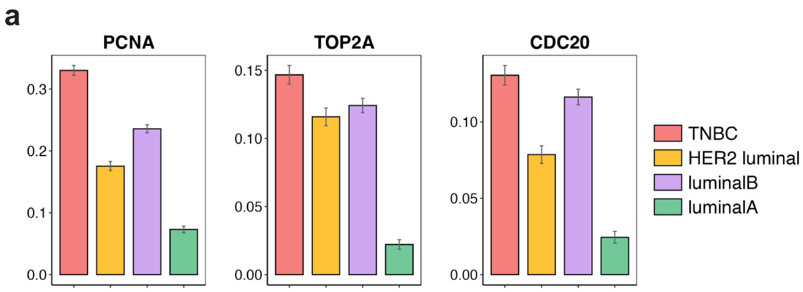

**Supplementary Fig. 9 | a.** Spatial annotations of the tissue regions in Section 1 and Section 2. **b.** Spatial protein expression patterns in Section 1 and Section 2.

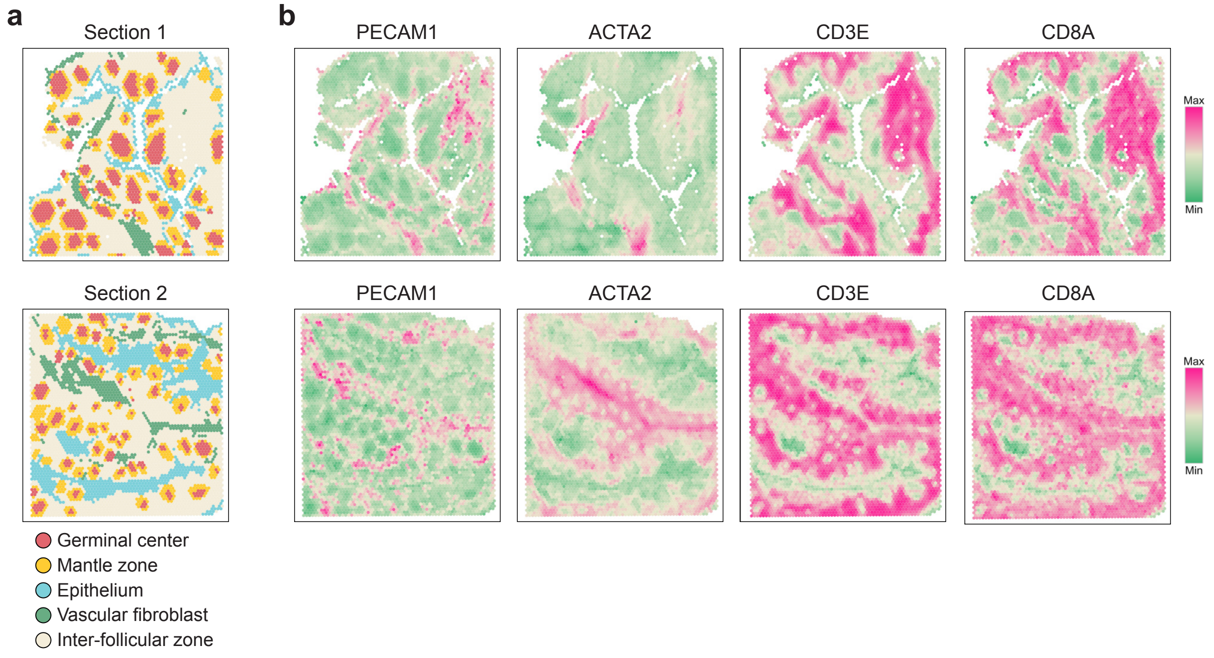

**Supplementary Fig. 10 | a.** Bar plots comparing region-specific F1 score under different modality settings (gene expression alone versus gene expression plus protein abundance) for supervised benchmark methods, demonstrating the effect of incorporating protein information on region-specific detection performance. SpaDo was excluded because it does not support protein modality input.

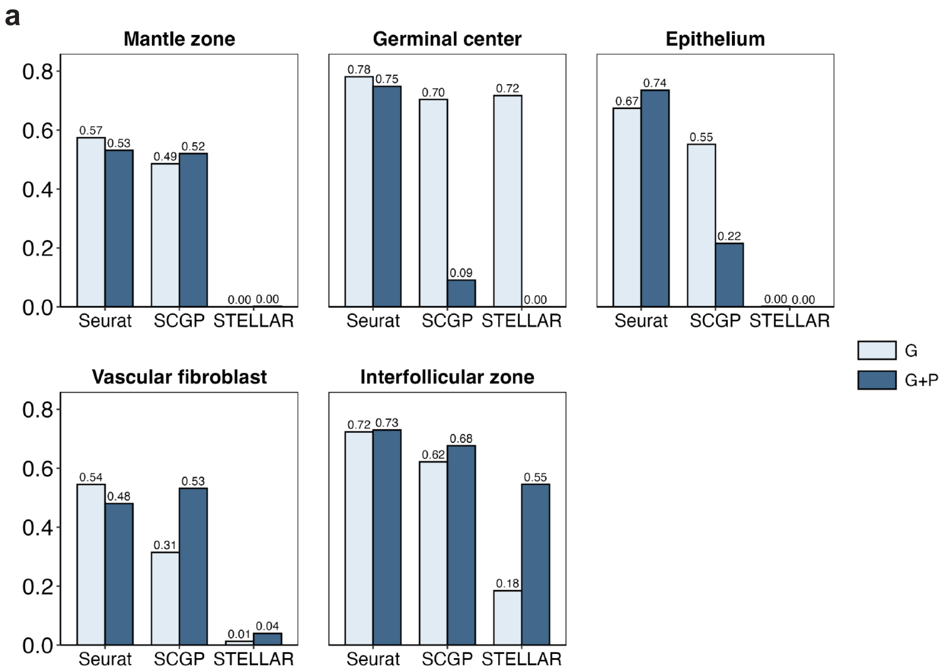

**Supplementary Fig. 11 | a.** Tissue region annotation results on mouse brain query samples.

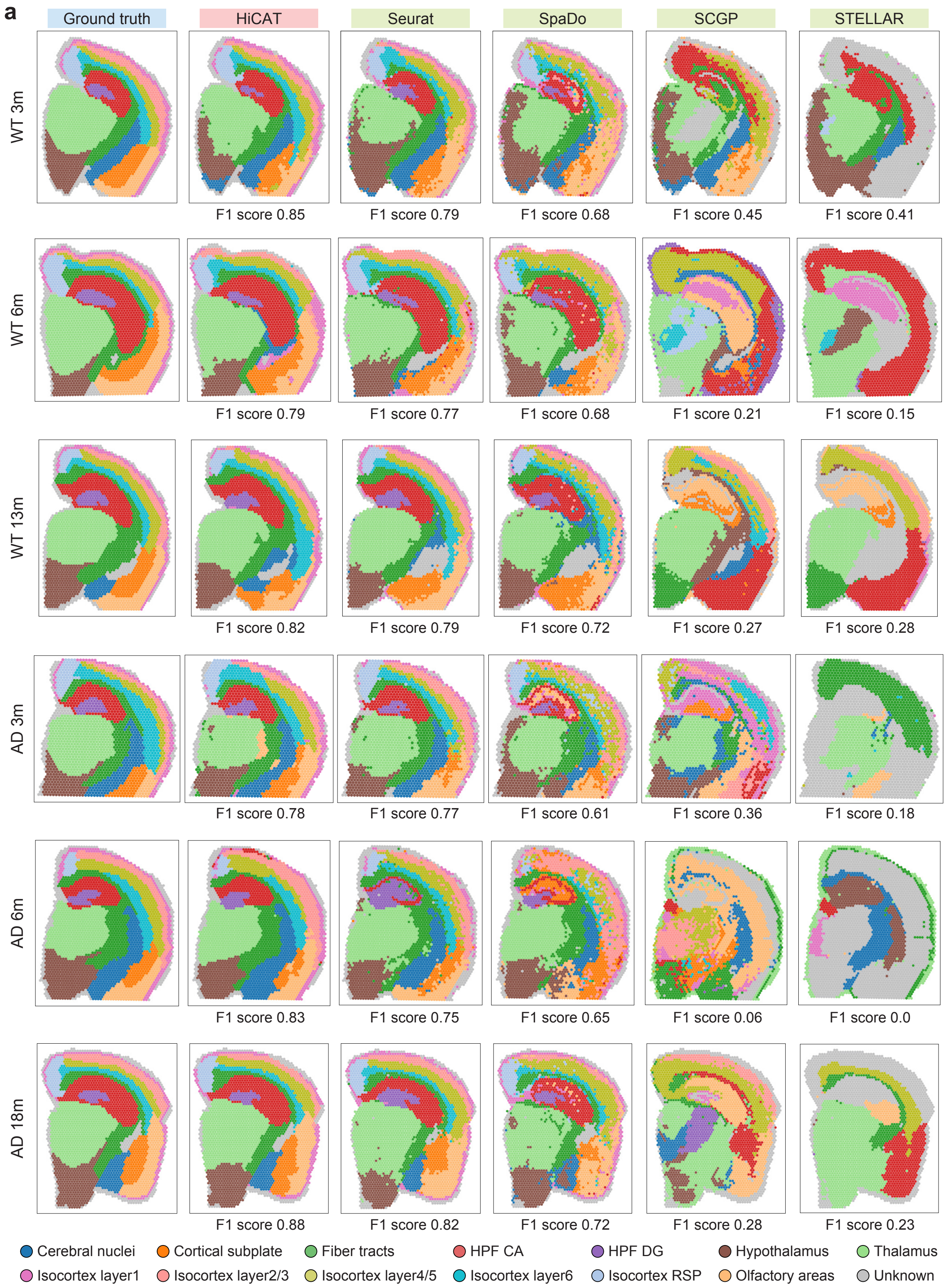

**Supplementary Note 5: Region-specific detection performances across supervised methods in a mouse brain dataset**

As shown in **Supplementary Fig. 12a** and **12b**, HiCAT achieved the highest region-specific precision and F1 score for most individual regions among all supervised methods. Compared with the second-best method, Seurat, HiCAT improved region-specific detection precision by an average of 5%, indicating an overall performance gain across regions. More importantly, detailed examination in **Supplementary Fig. 12c** showed that HiCAT provided stronger performance in resolving fine-scale brain structures. Within the hippocampal formation, HiCAT improved accuracy by 14%-1677% and F1 score by 13%-1073% relative to other methods when differentiating intricate substructures. Similarly, within the isocortex, HiCAT improved accuracy by 7%-815% and F1 score by 5%-927% for distinguishing cortical layers. These results highlight the effectiveness of HiCAT in resolving biologically meaningful fine-scale anatomical structures.

**Supplementary Fig. 12 |** Bar plots comparing region-specific **a.** precision and **b.** F1 score across supervised methods in mouse brain dataset. **c.** Bar plots comparing the performance of supervised label transfer methods in distinguishing substructures within the hippocampal formation and cortical layers within the isocortex. Bar heights represent the mean accuracy and F1 score across query samples, and error bars indicate standard errors.

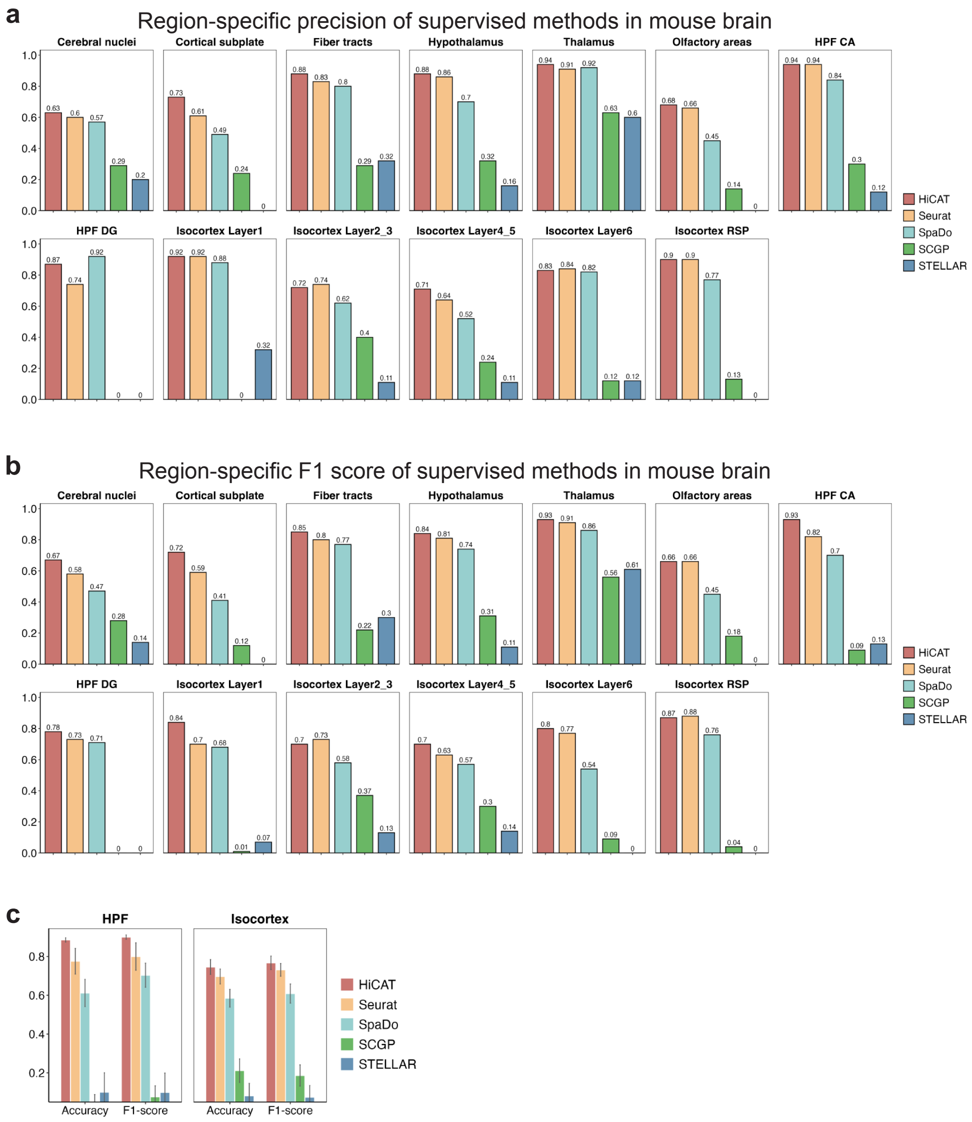

**Supplementary Note 6: HiCAT infers biologically meaningful hierarchical clonal tree structures in prostate cancer**

To evaluate whether HiCAT can infer biologically meaningful hierarchical structures, we applied the tree inference framework to prostate cancer 10x Visium datasets. In the original study[11], copy number status for each spot was inferred from spatially resolved mRNA profiles, and tumor spots were grouped into distinct clones. Based on the inferred copy number variation across clones, phylogenetic trees were constructed to represent the sequential order and similarity of clonal events. We used these published phylogenetic trees as ground truth to assess whether HiCAT could recover clonal relationships from multimodal spatial transcriptomics inputs.

We evaluated HiCAT under both single-section and multi-section settings. In the single-section analysis of section H2_1 (**Supplementary Fig. 13a-c**), the HiCAT-inferred binary hierarchy captured the relative similarity among tumor clones. By treating the shared latent clone structure shown in grey in the phylogenetic tree as a common branch node, the HiCAT-inferred tree achieved 71% similarity with the published ground truth. Although it is difficult to resolve the hierarchical relationships among clones A, C, and D in the phylogenetic tree, the HiCAT-inferred structure provides a plausible hierarchical interpretation of their developmental relationships. In the multi-section analysis of sections H1_4 and H1_5 (**Supplementary Fig. 13d-f**), HiCAT achieved 100% alignment with the published phylogenetic tree using the same evaluation strategy.

Together these results show that HiCAT can effectively leverage multimodal spatial transcriptomics data to infer biologically meaningful hierarchical structures. They also demonstrate the flexibility of the HiCAT tree inference framework, which can incorporate diverse label types, including pathologist annotations and clonal clusters, to support both label transfer and characterization of tumor clonal architecture.

**Supplementary Fig. 13 | a – c.** Spatial distribution and hierarchical relationships of tumor clones in section H2_1. **a.** Spatial visualization of tumor clones A – G. **b.** Phylogenetic clone tree of the tumor clones. **c.** HiCAT-inferred hierarchical clone tree structure.
**d – f.** Spatial distribution and hierarchical relationships of tumor clones in sections H1_4 and H1_5. **d.** Spatial visualization of tumor clones E – K. **e.** Phylogenetic clone tree of the tumor clones. **f.** HiCAT-inferred hierarchical clone tree structure.

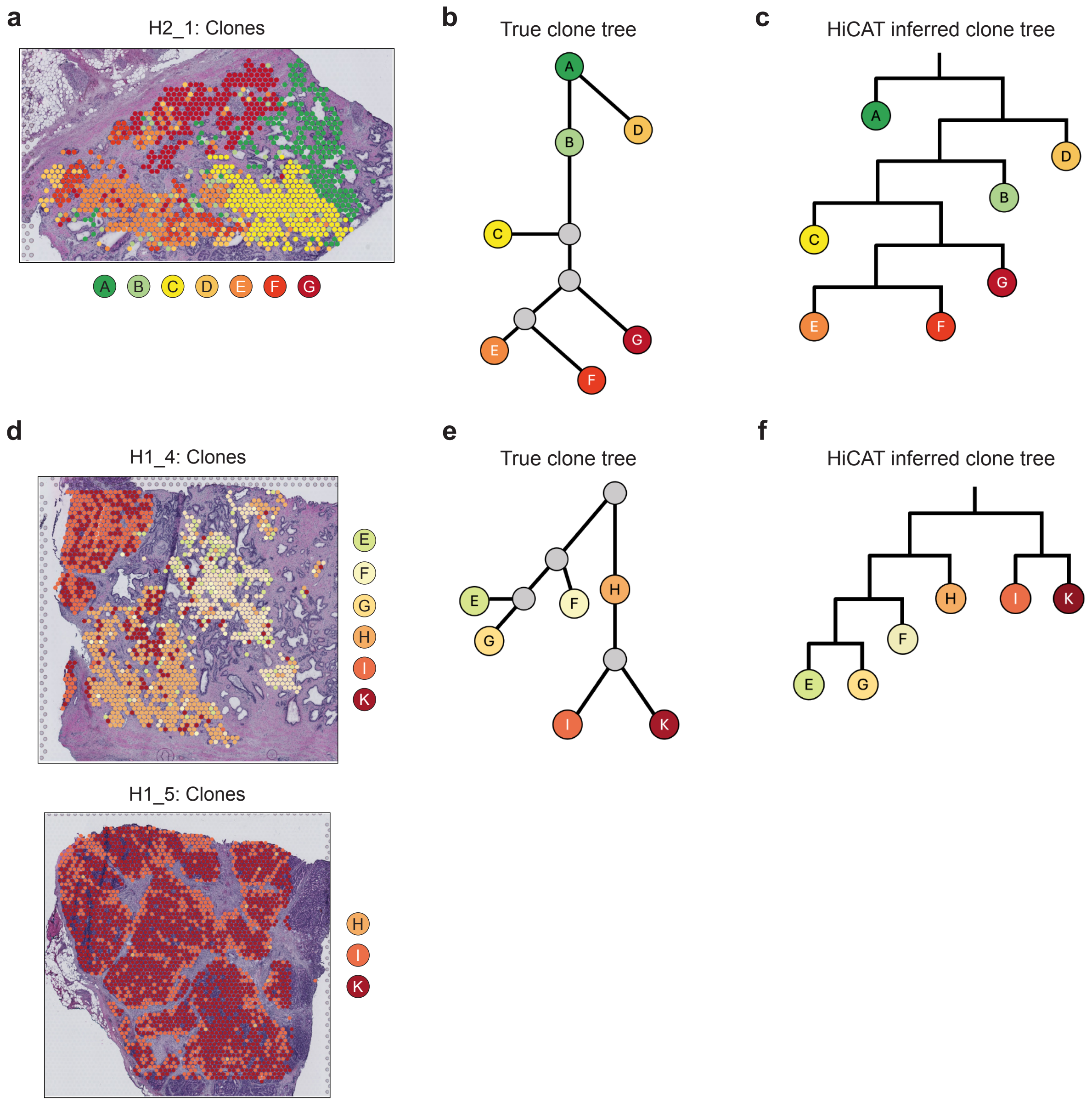
